# Linguistic structure and meaning organize neural oscillations into a content-specific hierarchy

**DOI:** 10.1101/2020.02.05.935676

**Authors:** Greta Kaufeld, Hans Rutger Bosker, Phillip M. Alday, Antje S. Meyer, Andrea E. Martin

**Author notes:** Greta Kaufeld; Max Planck Institute for Psycholinguistics, P.O Box 310, 6500AH Nijmegen, the Netherlands. The authors declare no competing financial interests.

## Abstract

Neural oscillations track linguistic information during speech comprehension (e.g., Ding et al., 2016; Keitel et al., 2018), and are known to be modulated by acoustic landmarks and speech intelligibility (e.g., Zoefel & VanRullen, 2015). But, it is unclear what information (e.g., timing, rhythm, or content) the brain utilizes to generate linguistic structure and meaning beyond the information that is present in the physical stimulus. We used electroencephalography (EEG) to investigate whether oscillations are modulated by linguistic content over and above the speech stimulus’ rhythmicity and temporal distribution. We manipulated the presence of semantic and syntactic information apart from the timescale of their occurrence, and controlled for the acoustic-prosodic and lexical-semantic information in the signal. EEG was recorded while 29 adult native speakers of all genders listened to naturally-spoken Dutch sentences, jabberwocky controls with a sentence-like prosodic rhythm and morphemes, word lists with lexical content but no phrase structure, and backwards acoustically-matched controls. Mutual information (MI) analysis revealed sensitivity to linguistic content: Phase MI was highest for sentences at the phrasal (0.8-1.1 Hz) and lexical timescale (1.9-2.8 Hz), suggesting that the delta-band is modulated by lexically-driven combinatorial processing beyond prosody, and that linguistic content (i.e., structure and meaning) organizes the phase of neural oscillations beyond the timescale and rhythmicity of the stimulus. This pattern is consistent with neurophysiologically-inspired models of language comprehension (Martin, 2016, 2020; Martin & Doumas, 2017) where oscillations encode endogenously-generated linguistic content over and above exogenous or stimulus-driven timing and rhythm information.

**Significance Statement:** Biological systems like the brain encode their environment not only by reacting in a series of stimulus-driven responses, but by combining stimulus-driven information with endogenous, internally-generated, inferential knowledge and meaning. Understanding language from speech is the human benchmark for this. Much research focusses on the purely stimulus-driven response, but here, we focus on the goal of language behavior: conveying structure and meaning. To that end, we use naturalistic stimuli that contrast acoustic-prosodic and lexical-semantic information to show that, during spoken language comprehension, oscillatory modulations reflect computations related to inferring structure and meaning from the acoustic signal. Our experiment provides the first evidence to date that compositional structure and meaning organize the oscillatory response, above and beyond acoustic and lexical controls.

## Introduction

There continues to be vibrant controversy about how the brain represents and processes biological signals that are highly relevant to humans, such as speech and language. How the brain maps the acoustics of speech onto abstract structure and meaning remains a core question across cognitive science and neuroscience. A large body of research has shown that neural populations closely align their phase to the envelope of the speech signal, which correlates with the syllable rate (e.g., Peelle & Davis, 2012; Zoefel & VanRullen, 2015; see Obleser & Kayser, 2019, for an excellent discussion of the use of the term “entrainment”). However, the degree to which neural responses encode *higher-level* linguistic information beyond the syllable level is relatively poorly understood. Delta-band oscillations have been linked to the top-down generation of hierarchically structured representations, such as phrases, clauses and sentences (e.g., Ding et al., 2017; Ding, Melloni, Zhang, Tian, & Poeppel, 2016; Meyer, Henry, Gaston, Schmuck, & Friederici, 2016), but these linguistic units do not exhibit a diagnostic physical manifestation in the acoustic signal in a straight-forward manner,^1^ so it is unclear what the brain “entrains” to. Here, we hypothesize that phase alignment (or “entrainment in the broad sense”, as defined by Obleser & Kayser, 2019, see also Haegens & Golumbic, 2018 and Meyer, Sun, & Martin, 2019) of neuronal populations to linguistic units that do *not* have straight-forward physically diagnostic representations in the acoustic signal may reflect a process of perceptual inference (Martin, 2016, 2020), whereby biological systems like the brain encode their environment not only by reacting in a series of stimulus-driven responses, but by combining stimulus-driven information with endogenous, internally-generated, inferential knowledge and meaning (Meyer, Sun, & Martin, 2019).

Results reported by Ding et al. (2016, 2017), Gross et al. (2013) and Keitel et al. (2018) yielded promising evidence that cortical networks may, indeed, track intrinsically generated syntactic structures beyond the syllable envelope. However, these previous experiments could not discriminate between cortical responses being driven by the *timing* or *frequency* of linguistic structures (cf. Frank & Yang, 2018), or by the *informational content of the structure itself.* We view this distinction as crucial: does the brain track inferred higher-level units such as phrases and words simply because they can be inferred at a given frequency, or does it attune to whether these units are structurally and semantically meaningful?

As a first step in understanding how listeners infer linguistic information from the acoustic signal, we ask if the neural response selectively attunes to stimuli as a function of our knowledge of language, and not just as a function of the timescales that information occurs on. In other words, is neural tracking of natural speech driven by structure and meaning beyond acoustic-prosodic modulations and lexical, non-compositional semantics?

Participants listened to naturally spoken, structurally homogenous sentences, jabberwocky items, and word lists (see Table 1). These three forward conditions were constructed in order to contrast two core sources of linguistic representations: prosodic structure, which can, but does not always, correlate with syntactic and information structure, and lexical semantics, which arises in isolated words and concepts. Additionally, we used backward speech as the core control of our experimental design, because it has an identical modulation spectrum for each forward condition. Using electroencephalography (EEG), we analyzed phase tracking of units at linguistically relevant timescales as quantified by Mutual Information (MI) – a typical measure of neural tracking that captures the informational similarity between two signals (Gross et al., 2013; Kayser, Ince, Gross, & Kayser, 2015; Keitel et al., 2018; Keitel, Ince, Gross, & Kayser, 2017). Figure 1 shows an overview of the experimental design and analysis pipeline. In sum, our experiment offers novel insights into how linguistic structure and meaning influence the neural response *above and beyond* the timescales at which that information occurs.

**Table 1.**
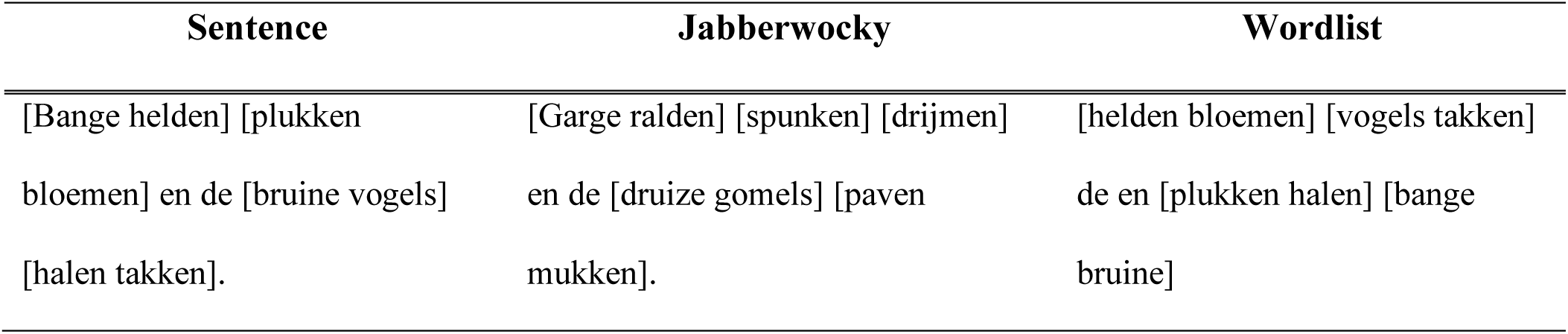
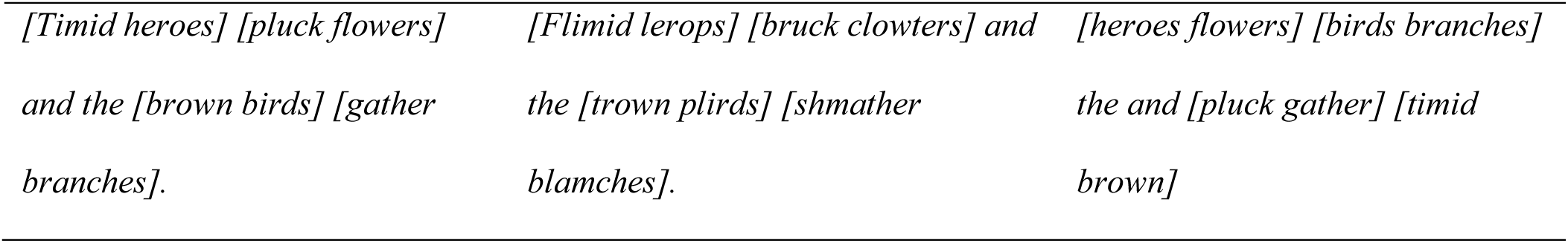
*Example items in Sentence, Jabberwocky, and Wordlist conditions.* Sentences consisted of 10 words (disyllabic, except for “de” (*“the”)* and “en” (*“and”*)) and carried sentence prosody. Jabberwocky items consisted of 10 pseudo-words with morphology; they also carried sentence prosody. Word lists consisted of the same 10 words as the Sentence condition, but scrambled so as to be syntactically implausible. They had list-prosody. Marked with square brackets are “phrases” in all three conditions.

**Figure 1.**
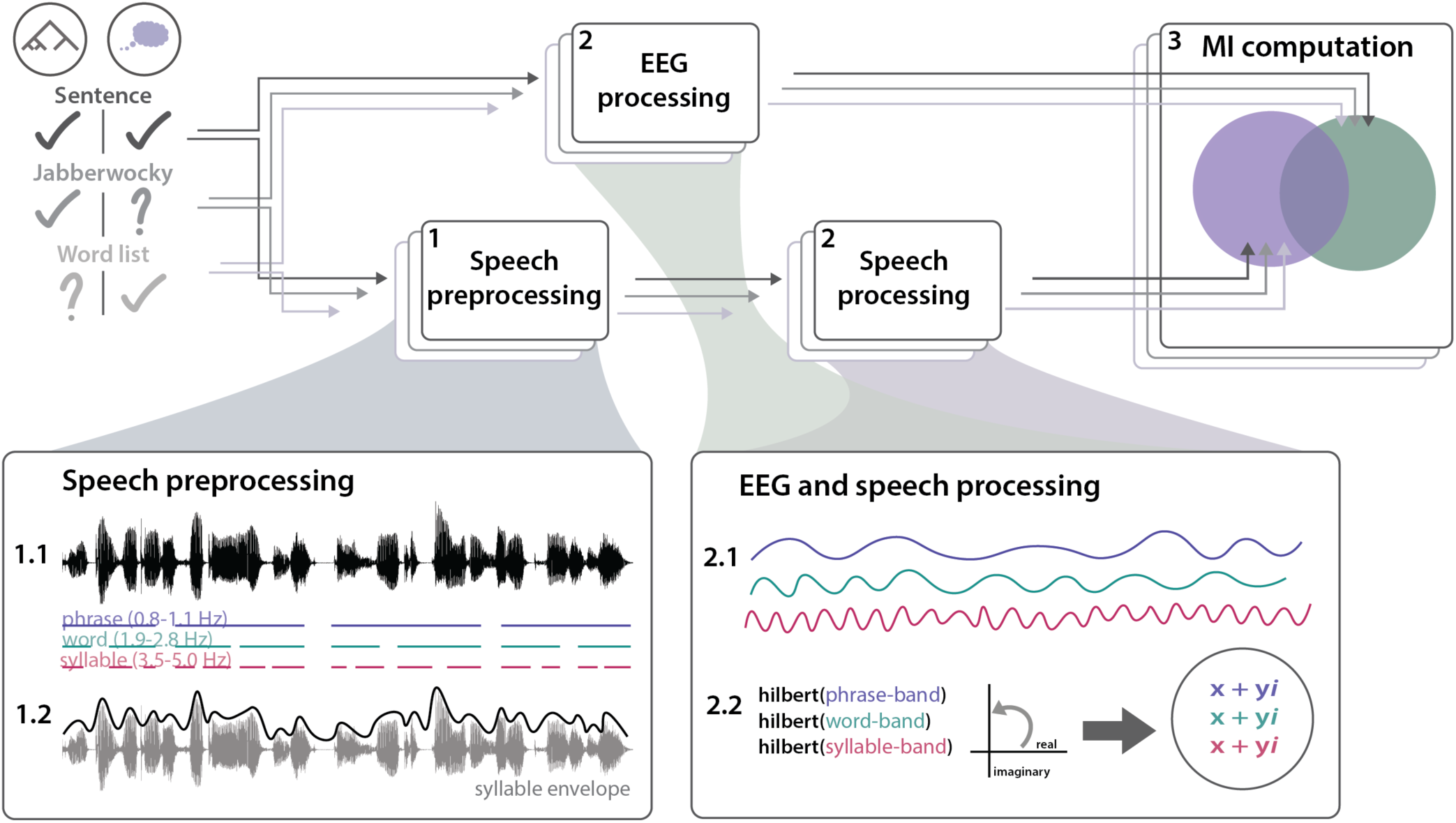
Experimental design and analysis pipeline. Participants listened to sentences, jabberwocky items, and word lists while their brain response was recorded using EEG. Step 1: Speech Processing. 1.1) The speech signal is annotated for the occurrence of phrases, words, and syllables in the stimuli. Based on this, frequency bands of interest for each of the linguistic units can be identified. 1.2) A cochlear filter is applied to the speech stimuli and the amplitude envelope is extracted. Step 2: Further processing is identical for both speech and EEG modalities. 2.1) Broadband filters are applied in the previously identified frequency bands of interest. 2.2) Hilbert transforms are computed in each filtered signal, and real and imaginary parts of the Hilbert transform output are used for further analysis. Step 3: MI Computation. Mutual information is computed between the pre-processed speech and EEG signal in each of the three conditions and their respective backward controls.

## Materials and Methods

### Participants

25 native Dutch speakers (9 males; age range 19-32; mean age=23) participated in the experiment. They were recruited from the Max Planck Institute for Psycholinguistics’ participant database with written consent approved by the Ethics Committee of the Social Sciences Department of Radboud University (Project code: ECSW2014-1003-196a). All participants in the experiment reported normal hearing and were remunerated for their participation.

### Materials

The experiment used three critical conditions: Sentence, Jabberwocky, and Wordlist. Eighty sets (triplets) of the three conditions (Sentence, Jabberwocky, Wordlist) were created, resulting in 240 stimuli. In addition to one “standard” forward presentation of each stimulus, participants also listened to a version of each of the stimuli played backwards, thus resulting in a total of 480 stimuli.

Dutch stimuli consisted of 10 words, which were all disyllabic except for “de” (*the*) and “en” (*and*), thus resulting in 18 syllables in total. Sentences all consisted of two coordinate clauses, which followed the structure *[Adj N V N Conj Det Adj N V N].* Word lists consisted of the same 10 words as in the Sentence condition, but scrambled in syntactically implausible ways (either *[V V Adj Adj Det Conj N N N N],* or *[N N N N Det Conj V V Adj Adj]*, in order to avoid any plausible internal combinations of words). Jabberwocky items were created using the wuggy pseudoword generator (Keuleers & Brysbaert, 2010), following the same syntactic structure as the Sentences. Table 1 shows an example of stimuli in each condition.

Forward stimuli were recorded by a female native speaker of Dutch in a sound-attenuating recording booth. All stimuli were recorded at a sampling rate of 44100 Hz (mono), using the Audacity sound recording and analysis software (Audacity Team, 2014). After recording, pauses were normalized to ∼150 ms in all stimuli, and the intensity was scaled to 70 dB using the Praat voice analysis software (Boersma & Weenink, 2017). Stimuli from all three conditions were then reversed using Praat. Figure 3 shows modulation spectra for forward and backward conditions. Please see the Appendix (Figure B and Tables 5a-c) for pilot data confirming participants’ perception of the linguistic content of each condition.

**Figure 2.**
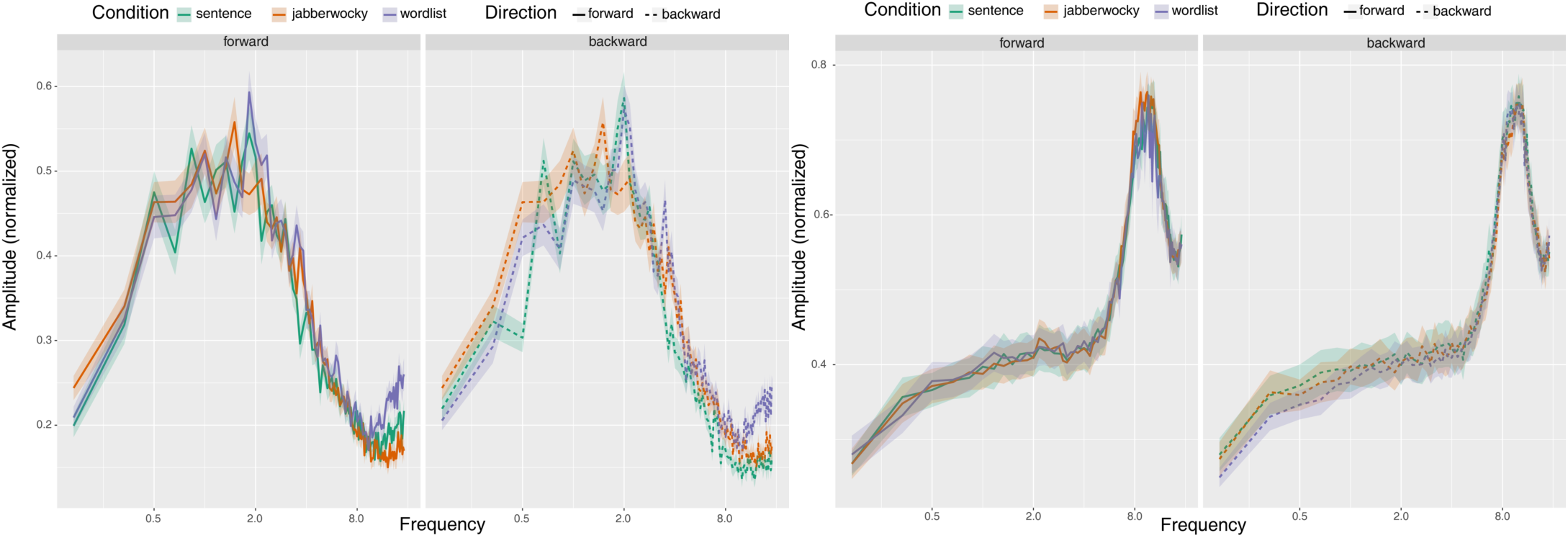
Modulation Spectra of forward and backward stimuli. Green: Sentences; Orange: Jabberwocky; Purple: Wordlist. Modulation spectra were calculated following the procedure and Matlab script described in Ding et al. (2017). Note that small deviations between the modulation spectrum of each forward condition and its backward counterpart are due to numerical inaccuracy; mathematically, the frequency components of forward and backward stimuli are identical.

**Figure 3.**
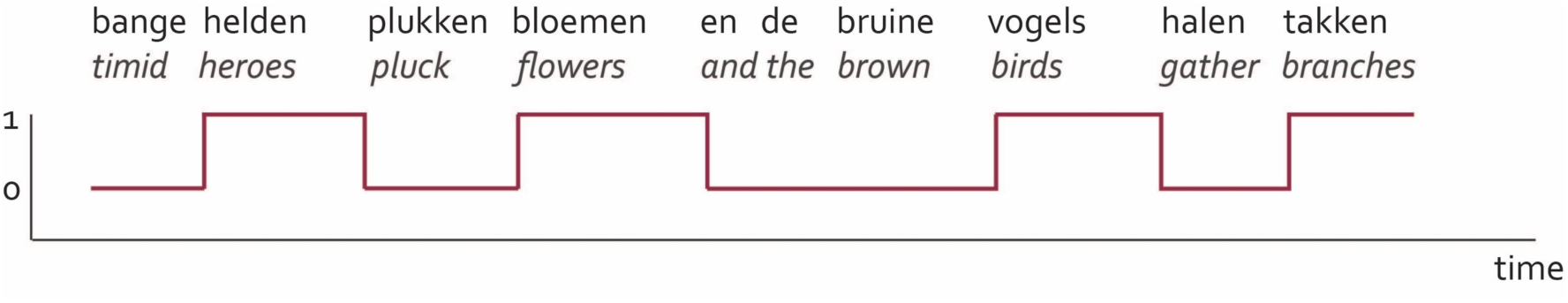
Visualization of the phrase-level annotations (inspired by Fig.2 in Brodbeck et al., 2018). Across time, the response array takes value 0 for words that cannot (yet) be integrated into phrases, and value 1 for words that can, resulting in a “pulse train” array.

### Procedure

Participants were tested individually in a sound-conditioned and Faraday-cage enclosed booth. They first completed a practice session with 4 trials (one from each forward condition and one backward example) to become familiarized with the experiment. All 80 stimuli from each condition were presented to the participants in separate blocks. The order of the blocks was pseudorandomized across listeners, and the order of the items within each block was randomized. During each trial, participants were instructed to look at a fixation cross which was displayed at the center of the screen (to minimize eye movements during the trial), and listen to the audio, which was presented to them at a comfortable level of loudness. After each trial, participants could advance to the next item via a button-press. After each block, participants were allowed to take a self-paced break. The experiment was run using the Presentation software (Neurobehavioral Systems) and took about 50-60 minutes to complete. EEG was continuously recorded with a 64-channel EEG system (MPI equidistant montage) using BrainVision Recorder^TM^ software, digitized at a sampling rate of 500 Hz and referenced to the left mastoid. The impedance of electrodes was kept below 25 kO. Data was re-referenced offline to the electrode average.

### EEG Data Preprocessing

The analysis steps were carried out using the FieldTrip analysis toolkit revision 20180320 (Oostenveld, Fries, Maris, & Schoeffelen, 2011) on MATLAB version 2016a (MathWorks, Inc.). Six participants were excluded from the analysis due to excessive artifact contamination. The raw EEG signal was segmented into a series of variable length epochs, starting at 200 ms before the onset of the utterance and lasting until 200 ms after its end. The signal was low-pass filtered to 70 Hz, and a 50-Hz bandpass filter was applied in each epoch to exclude line noise (both zero-phase FIR filters using Hamming windows). Channels contaminated with excessive noise were excluded from the analysis. Independent component analysis was performed on the remaining channels, and components related to eye movements, blinking, or motion artifacts, were subtracted from the signal. Epochs containing voltage fluctuations exceeding ±100 µV or transients exceeding ±75 µV were excluded from further analysis. We selected a cluster of 22 electrodes for all further analyses based on previous studies that found broadly-distributed effects related to sentence processing (e.g., Kutas, Van Petten & Kluender, 2006; Kutas & Federmeier, 2000; see also Ding et al., 2017). Specifically, the electrode selection included the following electrodes: 1, 2, 3, 4, 5, 8, 9, 10, 11, 28, 29, 30, 31, 33, 34, 35, 36, 37, 40, 41, 42, 43 (electrode names based on the MPI equidistant layout). We note that our results also hold for all electrodes (see Supplementary Materials).

### Speech Preprocessing

For each stimulus, we computed the wideband speech envelope at a sampling rate of 150 Hz following the procedure reported by Keitel et al. (2018) and others (e.g., Bosker & Cooke, 2018; Gross et al., 2013; Keitel et al., 2017). We first filtered the acoustic waveforms into 8 frequency bands (100-8,000 Hz; third-order Butterworth filter, forward and reverse), equidistant on the cochlear frequency map (Smith, Delgutte, & Oxenham, 2002). We then estimated the wideband speech envelope by computing the magnitude of the Hilbert transformed signal in each band and averaging across bands.

The timescales of interest for further Mutual Information analysis were identified in a similar fashion as described in Keitel et al. (2018). We first annotated the occurrence of linguistic units (phrases, words, and syllables) in the speech stimuli. Here, phrases were defined as noun-adjective-verb combinations (e.g., in the Sentence condition: “bange helden” – *timid heroes*; “plukken bloemen” – *pluck flowers,* and so on.; in the Jabberwocky condition: “garge ralden” – *flimid lerops* etc.; in the Wordlist condition, a “pseudo-phrase” corresponds to adjacent noun-noun, verb-verb and adjective-adjective pairs, e.g., “helden bloemen” – *heroes flowers*). Unit-specific bands of interest were then identified by converting each of the rates into frequency ranges across conditions. This resulted in the following bands: 0.8-1.1 Hz (phrases); 1.9-2.8 Hz (words); and 3.5-5.0 Hz (syllables).^2^

For an additional, exploratory annotation-based MI analysis (section Results, subsection *Tracking of abstract linguistic units*), we further created linguistically abstracted versions of our stimuli. Specifically, our aim was to create annotations that captured linguistic information at the phrase frequency entirely independent of the acoustic signal. Based on the word-level annotations of our stimuli, we created dimensionality-reduced arrays for further analysis (cf. the “Semantic composition” analyses reported by Brodbeck et al., 2019). Specifically, we identified all time points in the spoken materials where words could be integrated into phrases, and marked each of these words associated with phrase composition (e.g., in a sentence such as “bange helden plukken bloemen en de bruine vogels halen takken” (*timid heroes pluck flowers and the brown birds gather branches*), the words “helden” (*heroes*), “bloemen” (*flowers*), “vogels” (*birds*) and “takken” (*branches*) were marked). All these critical words were coded as 1 for their entire duration, while all other timepoints (samples) were marked as 0 (cf. Brodbeck et al., 2019). This resulted in an abstract “spike train” array of phrase-level structure building that is independent of the acoustic envelope. We repeated this procedure for all items individually in all three conditions, since our stimuli were naturally spoken and thus differed slightly in duration and time course. Note that, consequently, this “phrase-level composition array” is somewhat arbitrary for the Wordlist condition as there are, per definition, no phrases in a word list. We annotated “pseudo-phrases” the same way as shown in Table 1. The procedure is visualized in Figure 3.

### Mutual Information Analysis

We used Mutual Information (MI) in order to quantify the statistical dependency between the phase of speech envelopes and the EEG recordings according to the procedure described in Keitel et al. (2018; see also Gross et al., 2013; Kayser et al., 2015; Keitel et al., 2017). Based on the previously identified frequency bands of interest (see section “Speech preprocessing” above), we filtered both speech envelopes and EEG signals in each band (third-order Butterworth filter, forward and reverse). We then computed the Hilbert transform in each band and used the part of the Hilbert transform corresponding to the phase information (see Ince et al., 2017, for details) for further analysis. This resulted in two sets of two-dimensional variables (one for speech signals and one for EEG responses) in each condition (forward and backward; again, see Ince et al., 2017, for a more in-depth description). To take brain-stimulus lag into account, we computed MI at 5 different lags, ranging from 60 to 140 ms in steps of 20 ms, and to exclude auditory evoked responses we excluded the first 200 ms of each stimulus-signal pair. MI values from all five lags were averaged for subsequent statistical evaluation. We further concatenated all trials from speech and brain signals in order to increase the robustness of MI computation (Keitel et al., 2018).

### Statistical Analysis

In order to test whether the statistical dependency between the speech envelope and the EEG data as captured by MI was modulated by the linguistic structure and content of the stimulus, we compared phase MI values in all three frequency bands separately. Linear mixed models were fitted to the log-transformed, trimmed (5% on each end of the distribution) phase MI values in each frequency band using lme4 (Bates et al., 2015) in R (R Core Team, 2018). Models included main effects of Condition (three levels: Sentence, Wordlist, Jabberwocky) and Direction (two levels: Forward, Backward), as well as their interaction. All models included by-participant random intercepts and random slopes for the Condition * Direction interaction. For model coefficients, degrees of freedom were approximated using Satterthwaite’s method as implemented in the package lmerTest (Kuznetsova, Brockhoff & Christensen, 2017). We used treatment coding in all models, with Sentence being the reference level for Condition, and Forward the reference level for Direction. We then computed all pairwise comparisons within each direction using estimated marginal means (Tukey correction for multiple comparisons) with emmeans (Length, 2018) in R (i.e., comparing Sentence Forward to Jabberwocky Forward and Wordlist Forward, but never Sentence Forward to Jabberwocky Backward, because we had no hypotheses about these comparisons). See Supplemental Materials (Tables 1a-3b) for complete model outputs.

**Table 1a.** Mixed-effects logistic regression results for log-transformed, trimmed phase MI in the phrase frequency band. ^a^ Contrast coded (treatment coding with Sentence as reference level); ^b^Contrast coded (treatment coding with Forward as reference level).

|  | Estimate | SE | df | t | p | Variance | SD |
| --- | --- | --- | --- | --- | --- | --- | --- |
| *Fixed effects* |  |  |  |  |  |  |  |
| Intercept | -4.421 | 0.086 | 29.473 | -51.433 | < 0.01 |  |  |
| Condition[T.Jabberwocky]<sup>a</sup> | -0.497 | 0.097 | 29.411 | -5.139 | < 0.01 |  |  |
| Condition[T.Wordlist]<sup>a</sup> | -0.402 | 0.117 | 29.369 | -3.421 | < 0.01 |  |  |
| Direction[T.Backward]<sup>b</sup> | -0.805 | 0.106 | 29.731 | -7.626 | < 0.01 |  |  |
| Condition:Direction | 0.368 | 0.150 | 29.086 | 2.455 | 0.02 |  |  |
| [T.Jabberwocky]:[T.Backward] |  |  |  |  |  |  |  |
| Condition:Direction | 0.336 | 0.168 | 29.099 | 2.011 | 0.06 |  |  |
| [T.Wordlist]:[T.Backward] |  |  |  |  |  |  |  |
| *Random effects* |  |  |  |  |  |  |  |
| Intercept \| Participant |  |  |  |  |  | 0.170 | 0.412 |
| Condition[T.Jabberwocky]<sup>a</sup> \| Participant |  |  |  |  |  | 0.187 | 0.431 |
| Condition[T.Wordlist]<sup>a</sup> \| Participant |  |  |  |  |  | 0.315 | 0.561 |
| Direction[T.Backward]<sup>b</sup> \| Participant |  |  |  |  |  | 0.236 | 0.486 |
| Condition[T.Jabberwocky]<sup>a</sup>:Direction[T.Backward]<sup>b</sup> \| Participant |  |  |  |  |  | 0.479 | 0.692 |
| Condition[T.Wordlist]<sup>a</sup>:Direction[T.Backward]<sup>b</sup> \| Participant |  |  |  |  |  | 0.646 | 0.804 |

**Table 1b.** Pair-wise contrast results for log-transformed, trimmed phase MI per Direction in the phrase frequency band.

|  | Estimate | SE | df | t | *p* |
| --- | --- | --- | --- | --- | --- |
| *Forward* |  |  |  |  |  |
| Sentence – Jabberwocky | 0.497 | 0.099 | 29.97 | 5.049 | < 0.01 |
| Sentence – Wordlist | 0.402 | 0.120 | 29.99 | 3.361 | < 0.01 |
| Jabberwocky – Wordlist | -0.096 | 0.103 | 30.09 | -0.929 | 0.63 |
| *Backward* |  |  |  |  |  |
| Sentence – Jabberwocky | 0.130 | 0.122 | 30.00 | 1.060 | 0.55 |
| Sentence – Wordlist | 0.066 | 0.130 | 30.02 | 0.509 | 0.87 |
| Jabberwocky – Wordlist | -0.063 | 0.135 | 30.02 | -0.470 | 0.89 |

**Table 2a.** Mixed-effects logistic regression results for log-transformed, trimmed phase MI in the word frequency band. ^a^ Contrast coded (treatment coding with Sentence as reference level); ^b^Contrast coded (treatment coding with Forward as reference level).

|  | Estimate | SE | df | t | p | Variance | SD |
| --- | --- | --- | --- | --- | --- | --- | --- |
| *Fixed effects* |  |  |  |  |  |  |  |
| Intercept | -5.322 | 0.767 | 28.437 | -69.359 | < 0.01 |  |  |
| Condition[T.Jabberwocky]<sup>a</sup> | -0.380 | 0.121 | 28.790 | -3.143 | < 0.01 |  |  |
| Condition[T.Wordlist]<sup>a</sup> | -0.120 | 0.072 | 30.047 | -1.666 | 0.11 |  |  |
| Direction[T.Backward]<sup>b</sup> | -0.474 | 0.126 | 29.034 | -3.754 | < 0.01 |  |  |
| Condition:Direction | 0.174 | 0.176 | 29.124 | 0.989 | 0.33 |  |  |
| [T.Jabberwocky]:[T.Backward] |  |  |  |  |  |  |  |
| Condition:Direction | 0.057 | 0.142 | 29.074 | 0.404 | 0.69 |  |  |
| [T.Wordlist]:[T.Backward] |  |  |  |  |  |  |  |
| *Random effects* |  |  |  |  |  |  |  |
| Intercept \| Participant |  |  |  |  |  | 0.127 | 0.357 |
| Condition[T.Jabberwocky]<sup>a</sup> \| Participant |  |  |  |  |  | 0.338 | 0.581 |
| Condition[T.Wordlist]<sup>a</sup> \| Participant |  |  |  |  |  | 0.063 | 0.250 |
| Direction[T.Backward]<sup>b</sup> \| Participant |  |  |  |  |  | 0.378 | 0.615 |
| Condition[T.Jabberwocky]:Direction[T.Backward] \| Participant |  |  |  |  |  | 0.726 | 0.852 |
| Condition[T.Wordlist]:Direction[T.Backward] \| Participant |  |  |  |  |  | 0.412 | 0.642 |

**Table 2b.** Pair-wise contrast results for log-transformed, trimmed phase MI per Direction in the word frequency band.

|  | Estimate | SE | df | t | p |
| --- | --- | --- | --- | --- | --- |
| *Forward* |  |  |  |  |  |
| Sentence – Jabberwocky | 0.380 | 0.123 | 30.05 | 3.086 | 0.01 |
| Sentence – Wordlist | 0.120 | 0.073 | 29.72 | 1.633 | 0.25 |
| Jabberwocky – Wordlist | -0.260 | 0.140 | 30.07 | -1.858 | 0.17 |
| *Backward* |  |  |  |  |  |
| Sentence – Jabberwocky | 0.206 | 0.119 | 30.02 | 0.536 | 0.85 |
| Sentence – Wordlist | 0.062 | 0.117 | 30.02 | 0.536 | 0.854 |
| Jabberwocky – Wordlist | -0.144 | 0.102 | 30.04 | -1.415 | 0.35 |

**Table 3a.**
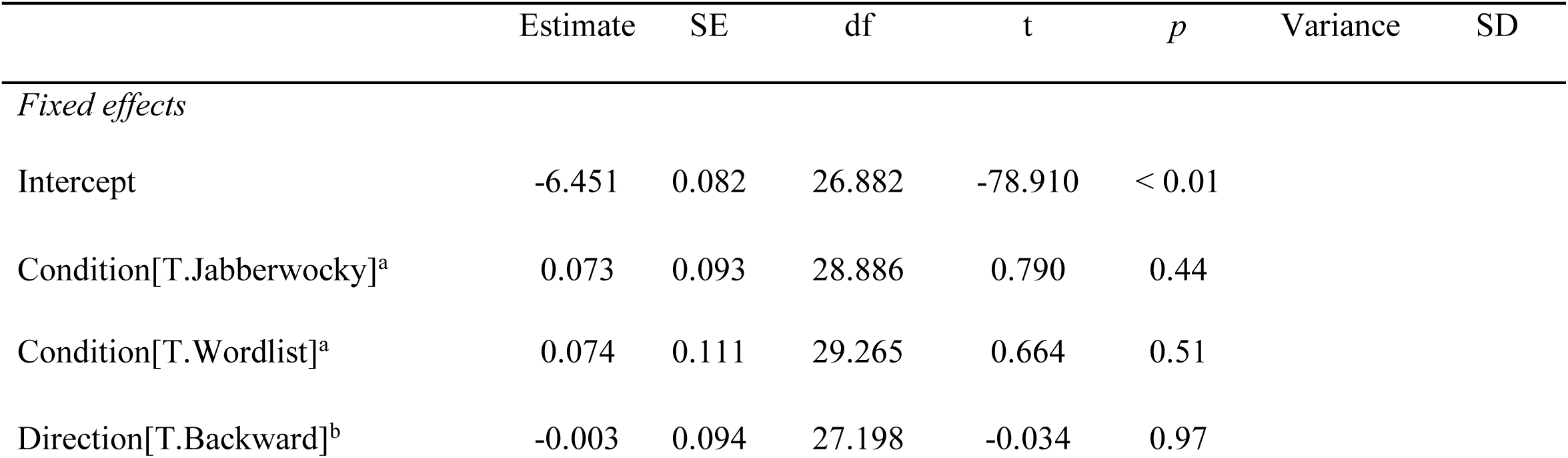
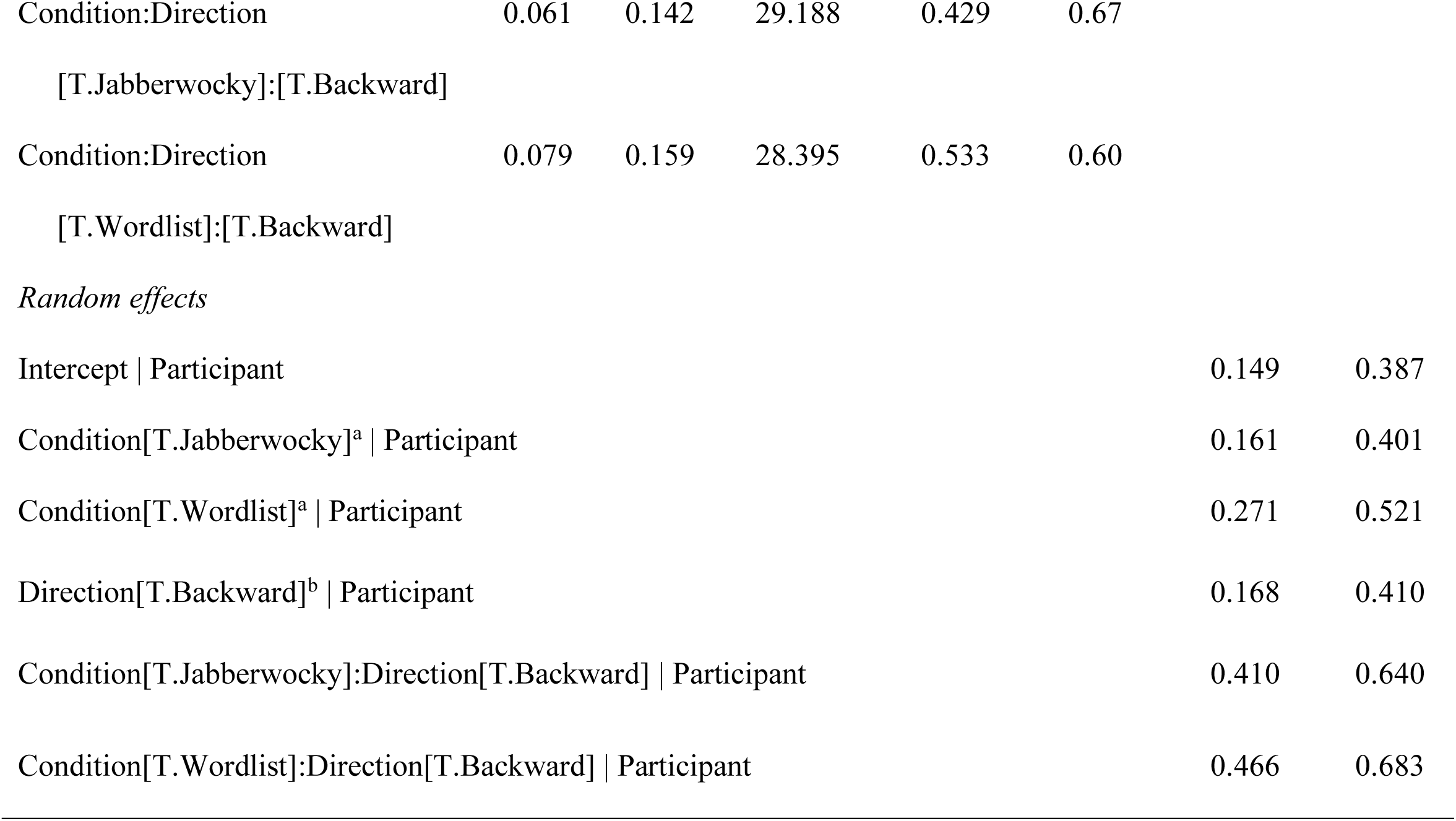
Mixed-effects logistic regression results for log-transformed, trimmed phase MI in the syllable frequency band. ^a^ Contrast coded (treatment coding with Sentence as reference level); ^b^Contrast coded (treatment coding with Forward as reference level).

|  | Estimate | SE | df | t | p | Variance | SD |
| --- | --- | --- | --- | --- | --- | --- | --- |
| *Fixed effects* |  |  |  |  |  |  |  |
| Intercept | -6.451 | 0.082 | 26.882 | -78.910 | < 0.01 |  |  |
| Condition[T.Jabberwocky]<sup>a</sup> | 0.073 | 0.093 | 28.886 | 0.790 | 0.44 |  |  |
| Condition[T.Wordlist]<sup>a</sup> | 0.074 | 0.111 | 29.265 | 0.664 | 0.51 |  |  |
| Direction[T.Backward]<sup>b</sup> | -0.003 | 0.094 | 27.198 | -0.034 | 0.97 |  |  |

|  |  |  |  |  |  |
| --- | --- | --- | --- | --- | --- |
| Condition:Direction | 0.061 | 0.142 | 29.188 | 0.429 | 0.67 |
| [T.Jabberwocky]:[T.Backward] |  |  |  |  |  |
| Condition:Direction | 0.079 | 0.159 | 28.395 | 0.533 | 0.60 |
| [T.Wordlist]:[T.Backward] |  |  |  |  |  |

*Random effects*
|  |  |  |
| --- | --- | --- |
| Intercept \| Participant | 0.149 | 0.387 |
| Condition[T.Jabberwocky]<sup>a</sup> \| Participant | 0.161 | 0.401 |
| Condition[T.Wordlist]<sup>a</sup> \| Participant | 0.271 | 0.521 |
| Direction[T.Backward]<sup>b</sup> \| Participant | 0.168 | 0.410 |
| Condition[T.Jabberwocky]:Direction[T.Backward] \| Participant | 0.410 | 0.640 |
| Condition[T.Wordlist]:Direction[T.Backward] \| Participant | 0.466 | 0.683 |

**Table 3b.**
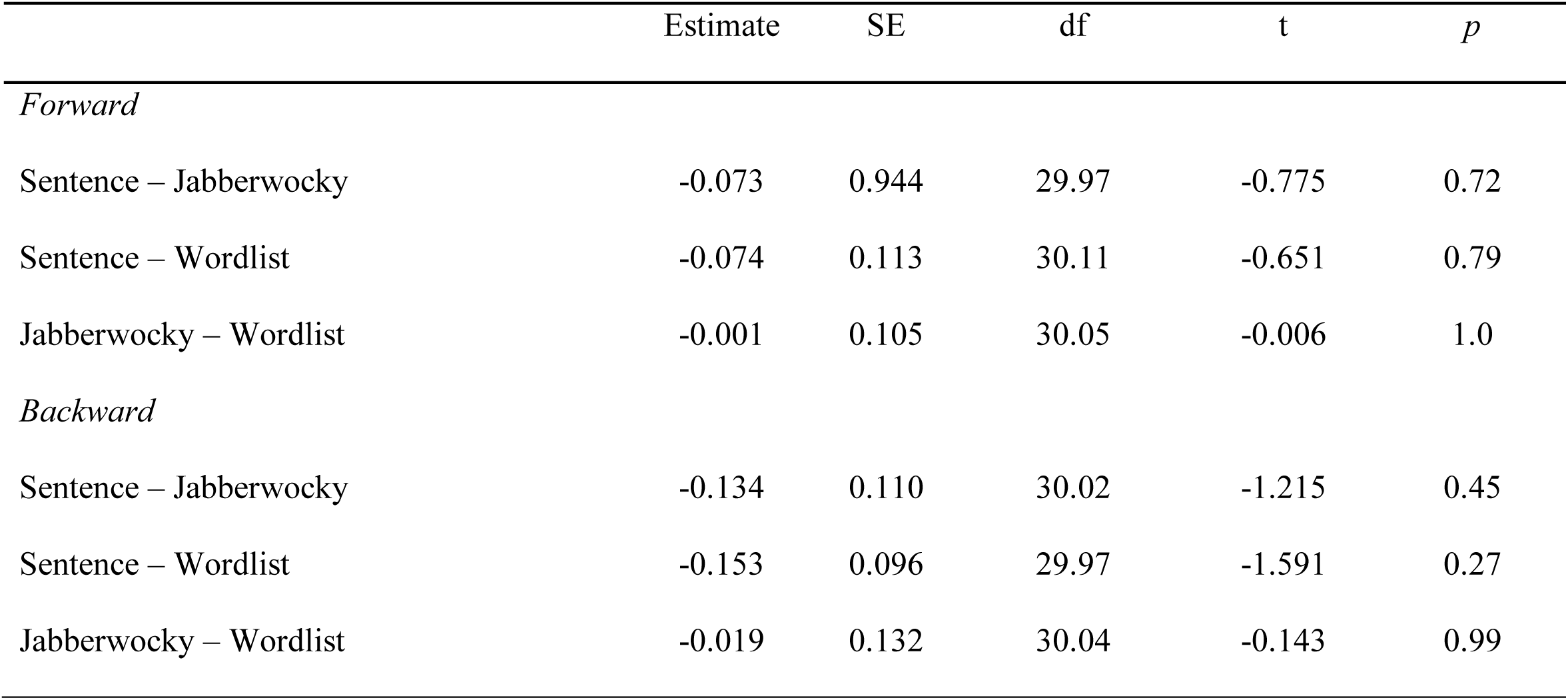
Pair-wise contrast results for log-transformed, trimmed phase MI per Direction in the syllable frequency band.

**Table 4a.** Mixed-effects logistic regression results for log-transformed, trimmed phase MI between brain and dimensionality-reduced stimuli. ^a^ Contrast coded (treatment coding with Sentence as reference level); ^b^Contrast coded (treatment coding with Forward as reference level).

|  | Estimate | SE | df | t | p | Variance | SD |
| --- | --- | --- | --- | --- | --- | --- | --- |
| *Fixed effects* |  |  |  |  |  |  |  |
| Intercept | -5.977 | 0.076 | 25.757 | -78.813 | < 0.01 |  |  |
| Condition[T.Jabberwocky]<sup>a</sup> | -0.328 | 0.095 | 29.219 | -3.436 | < 0.01 |  |  |
| Condition[T.Wordlist]<sup>a</sup> | -0.522 | 0.110 | 27.529 | -4.737 | < 0.01 |  |  |
| Direction[T.Backward]<sup>b</sup> | -0.685 | 0.112 | 25.071 | -6.135 | < 0.01 |  |  |
| Condition:Direction | 0.254 | 0.156 | 28.115 | 1.627 | 0.11 |  |  |
| [T.Jabberwocky]:[T.Backward] |  |  |  |  |  |  |  |
| Condition:Direction | 0.580 | 0.154 | 28.464 | 3.785 | < 0.01 |  |  |
| [T.Wordlist]:[T.Backward] |  |  |  |  |  |  |  |
| *Random effects* |  |  |  |  |  |  |  |
| Intercept \| Participant |  |  |  |  |  | 0.130 | 0.360 |
| Condition[T.Jabberwocky] \| Participant |  |  |  |  |  | 0.195 | 0.441 |
| Condition[T.Wordlist] \| Participant |  |  |  |  |  | 0.283 | 0.532 |
| Direction[T.Backward] \| Participant |  |  |  |  |  | 0.291 | 0.540 |
| Condition[T.Jabberwocky]:Direction[T.Backward] \| Participant |  |  |  |  |  | 0.572 | 0.756 |
| Condition[T.Wordlist]:Direction[T.Backward] \| Participant |  |  |  |  |  | 0.555 | 0.745 |

**Table 4b.**
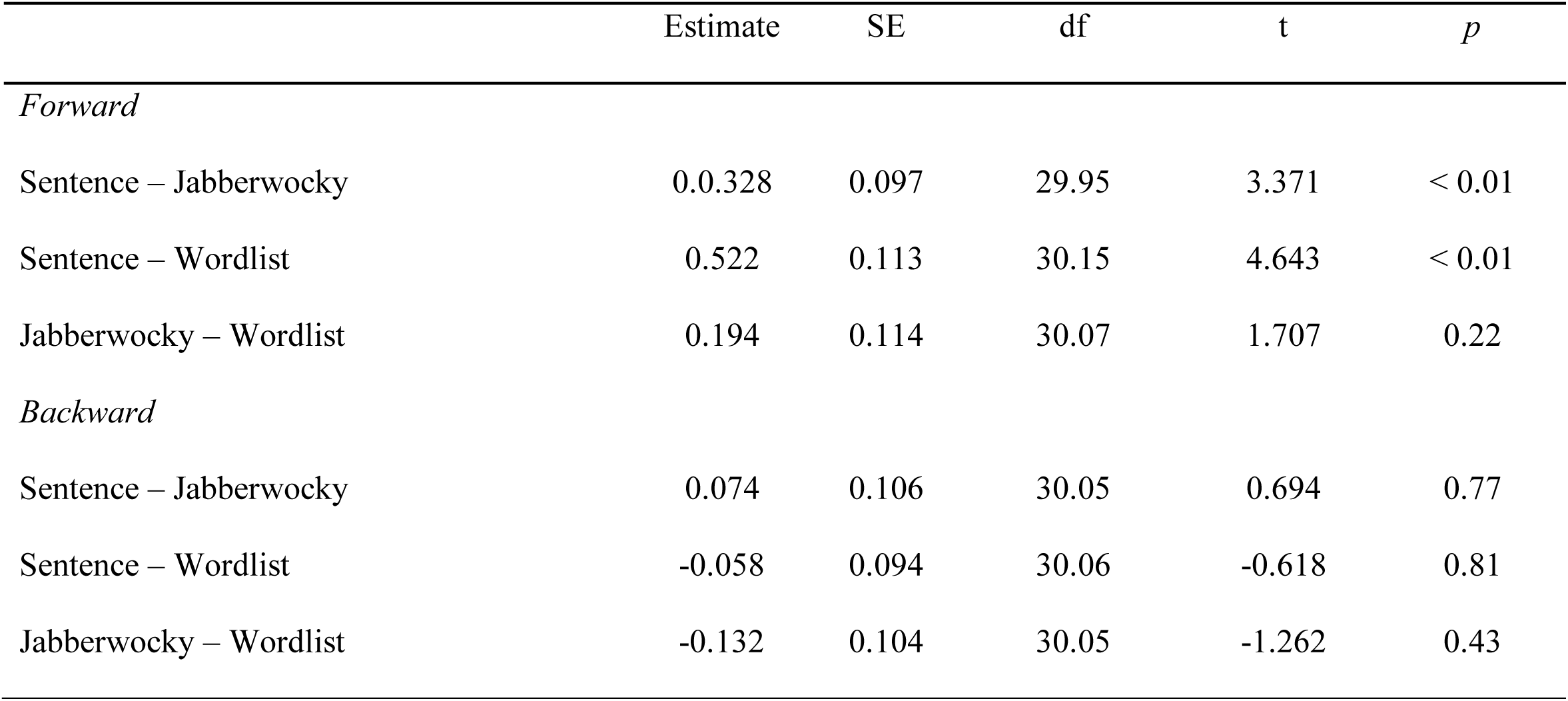
Pair-wise contrast results for log-transformed, trimmed phase MI between brain and dimensionality-reduced stimuli.

**Table 5a.** String token sort ratio (averaged across items) for Sentence, Jabberwocky and Wordlist items. Participants (N=7) listened to forward recordings of all items and were asked to type what they heard. String token sort ratios were computed using fuzzywuzzy in python.

| *Sentence* | *Jabberwocky* | *Wordlist* |
| --- | --- | --- |
| 94.96 | 32.37 | 72.42 |

**Table 5b.** Linear mixed model results for string token sort ratio during the pretest. ^a^ Contrast coded (treatment coding with Sentence as reference level). Model structure: Response ∼ Condition + (1|Item) + (1|Participant).

|  | Estimate | SE | t | Variance | SD |
| --- | --- | --- | --- | --- | --- |
| *Fixed effects* |  |  |  |  |  |
| Intercept | 94.96 | 2.02 | 47.14 |  |  |
| Condition[T.Jabberwocky]<sup>a</sup> | -62.59 | 0.88 | -71.28 |  |  |
| Condition[T.Wordlist]<sup>a</sup> | -22.54 | 0.88 | -25.67 |  |  |
| *Random effects* |  |  |  |  |  |
| Intercept \| Item |  |  |  | 8.57 | 2.93 |
| Intercept \| Participant |  |  |  | 24.94 | 4.99 |

**Table 5c.** Pair-wise contrast results for string token sort ratios during the pretest.

|  | Estimate | SE | df | t | p |
| --- | --- | --- | --- | --- | --- |
| *Forward* |  |  |  |  |  |
| Sentence – Jabberwocky | 62.59 | 0.88 | 1552 | 71.28 | < 0.01 |
| Sentence – Wordlist | 22.54 | 0.88 | 1552 | 25.67 | < 0.01 |
| Jabberwocky – Wordlist | -40.05 | 0.88 | 1552 | -45.61 | < 0.1 |

For the exploratory dimensionality-reduced MI analysis, we performed the same set of statistical analyses (but only in one single frequency band). Specifically, we fitted a linear mixed model including main effects of Condition (three levels: Sentence, Wordlist, Jabberwocky) and Direction (two levels: Forward, Backward), as well as their interaction and by-participant random intercepts and random slopes for the Condition * Direction interaction to the log-transformed, trimmed phase MI values. We then computed estimated marginal means precisely as described in the previous section.

Finally, in order to analyze whether phase MI was statistically higher at lower frequency bands, corresponding to more structured linguistic representations, we repeated the statistical analysis described above, but this time submitting the phase MI values from all three frequencies to a single mixed model that included fixed effects of Direction and Frequency, as well as their interaction and random by-participant intercepts. Again, degrees of freedom were approximated using Satterthwaite’s method as implemented in the package lmerTest (Kuznetsova, Brockhoff & Christensen, 2017). We used treatment coding in the model, with the syllable level being the reference level for Frequency, and Forward the reference level for Direction. We then computed all pairwise comparisons within each direction using estimated marginal means (Tukey correction for multiple comparisons) with emmeans in R as previously (Length, 2018).

## Results

### Speech tracking

We computed Mutual Information (MI) between the phase of the Hilbert-transformed EEG time series and the phase of the Hilbert-transformed speech envelope within three frequency bands of interest that corresponded to the occurrence rates of phrases (0.8-1.1 Hz), words (1.9-2.8 Hz), and syllables (3.5-5.0 Hz) in a cluster of central electrodes.

Specifically, we designed our experiment to assess whether the brain response is driven by the (quasi-)periodic temporal occurrence of linguistic structures, or whether it is modulated as a function of the *content* of those structures. Using MI allowed us to quantify and compare the degree of speech tracking across sentences, word lists, and jabberwocky items.

We found condition-dependent enhanced phase MI at distinct timescales (see Figure 2). In the phrase frequency band (0.8-1.1 Hz), phase MI was significantly higher for sentences compared to jabberwocky items and word list (Sentence – Jabberwocky: Δ = 0.497, SE = 0.099, *p* = 0.001; Sentence – Wordlist: Δ = 0.402, SE = 0.120, *p* = 0.006; all results Tukey corrected for multiple comparisons); in the word frequency band (1.9-2.8 Hz), phase MI was significantly higher for sentences compared to jabberwocky items (Sentence – Jabberwocky: Δ = 0.380, SE = 0.123, *p* = 0.012). None of these effects were present in the backward speech, demonstrating that they were not driven by the acoustic properties of the stimuli (see section Materials and Methods for details about our mixed effects models structures; see Supplemental Materials – Tables 1a, 1b, 2a and 2b – for complete model outputs). Note also that the likelihood ratio tests (comparable to the interaction term in classical repeated-measures Anova) revealed a significant interaction between Condition and Direction in the phrase frequency band (χ^2^(2) = 6.253, *p* = 0.044); on the word timescale, the pair-wise difference between the forward conditions was a simple effect (χ^2^(2) = 0.983, *p* = 0.612). An almost identical pattern of results also emerged when computing MI over all electrodes (rather than a cluster of central ones), with sentences showing higher phase MI than jabberwocky items and word lists both in the phrase band (Sentence – Jabberwocky: Δ = 0.356, SE = 0.076, *p* < 0.001; Sentence – Wordlist: Δ = 0.309, SE = 0.091, *p* = 0.005) and in the word frequency band (Sentence – Jabberwocky: Δ = 0.329, SE = 0.091, *p* = 0.003; Sentence – Wordlist: Δ = 0.139, SE = 0.046, *p* = 0.014). Again, we found no significant differences between the backward controls when computing MI over the entire head. These findings indicate that neural tracking is enhanced for linguistic structures at timescales specific to that structure’s role in the unfolding meaning of the sentence, consistent with neurophysiologically inspired models of language comprehension (Martin, 2016, 2020; Martin & Doumas, 2017).

Like previous studies (e.g., Gross et al. 2013; Kayser et al., 2015; Keitel et al., 2018), we also found significantly higher phase MI in lower frequency bands, corresponding to higher-level linguistic units such as phrases, compared to the word and syllable frequency bands (see Figure 2; phrase – word: Δ = 0.757, SE = 0.079, *p* < 0.001; phrase – syllable: Δ = 1.664, SE = 0.083, *p* < 0.001 word – syllable: Δ = 0.908, SE = 0.083, *p* < 0.001). These results might suggest that the brain response is largely driven by information present at lower frequency bands, rather than tracking of the syllable envelope. Note, however, that we found the same significant effects for comparisons between the backward conditions (phrase – word: Δ = 0.602, SE = 0.080, *p* < 0.001; phrase – syllable: Δ = 1.076, SE = 0.083, *p* < 0.001; word – syllable: Δ = 0.474, SE = 0.076, *p* < 0.001). Although the effects appear to be more pronounced for forward than for backward conditions, as evidenced by a significant positive Frequency * Direction interaction in the base model (χ^2^(2) = 26.293, *p* < 0.001; based on likelihood ratios), it is unclear whether increased MI at lower frequencies reflects increased sensitivity of the brain to information occurring at those frequencies, or whether these effects are due to properties of the acoustic signal and/or related to the 1/f frequency component (see Ouyang et al., 2019, for a more detailed discussion of the potential confounds of 1/f brain activity).

**Figure 4.**
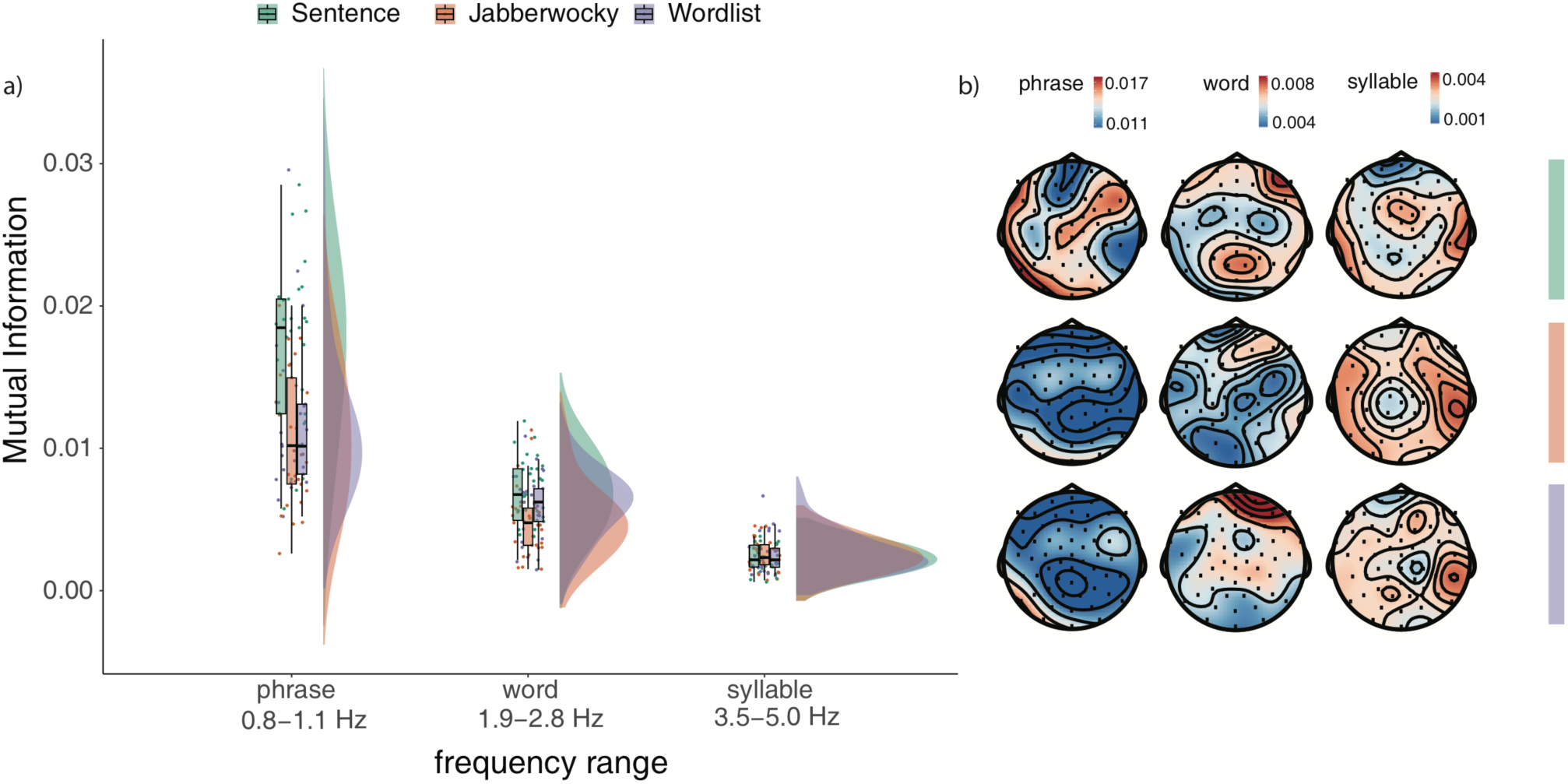
Phase MI between speech signal and brain response. Panel a) shows MI for Sentences (green), Jabberwocky items (orange), and Word lists (purple) for phrase, word and syllable timescales across central electrodes (each dot represents one participant’s mean MI response averaged across electrodes). Panel b) shows the average scalp distribution of MI per condition and band, averaged across participants. Raincloud plots were made using the Raincloud package in R (Allen et al., 2019).

### Tracking of abstract linguistic units

Inspecting the modulation spectra of our stimuli (Figure 2), it is apparent that – although carefully designed – the acoustic signals are not entirely indistinguishable between conditions based on their spectral properties. Most notably, Sentence stimuli appear to exhibit a small peak at around 0.5 Hz (roughly corresponding to the phrase timescale in our stimuli) compared to the other two conditions. It is important to note that (1) differences between conditions are not surprising, given that our stimuli were naturally spoken, and (2) we specifically designed our experiment to include backward versions of all conditions to control for slight differences between the acoustic envelopes of the forward stimuli. That being said, we conducted an additional, exploratory analysis of the phrase frequency band in order to further reduce the potential confound of differences between the acoustic modulation spectra, and to disentangle the distribution of linguistic phrase representations and the acoustic stimulus even further.^3^ Specifically, we computed phase MI in the delta-theta range (0.8-5 Hz) between the brain response and abstracted, dimensionality-reduced annotations of all stimuli, containing only information about when words could be integrated into phrases (cf. Brodbeck, Presacco & Simon, 2018; see section Materials and Methods for detailed descriptions of how these annotations were created). These annotation-based analyses revealed increased phase MI for sentences compared to jabberwocky items and word lists (Sentence – Jabberwocky: Δ = 0.328, SE = 0.097, *p* = 0.006; Sentence – Word list: Δ = 0.522, SE = 0.113, *p* = 0.002; all results Tukey corrected for multiple comparisons) and no significant difference among the backward controls (see Supplemental Materials for complete model outputs). Again, the same pattern of results also emerged when computing phase MI over all electrodes (Sentence – Jabberwocky: Δ = 0.354, SE = 0.076, *p* < 0.001; Sentence – Word list: Δ = 0.616, SE = 0.110, *p* < 0.001). These results support our previously reported findings, showing that phase alignment is influenced by the presence of abstract linguistic information. In other words, this exploratory analysis supports our earlier finding that the brain’s “sensitivity” to linguistic structure and meaning goes above and beyond the acoustic signal.

## Discussion

The current experiment tested how the brain attunes to inferred linguistic information that is not encoded in the acoustic properties of the stimulus in a straight-forward manner. Contrasting sentences, word lists and jabberwocky items, we analyzed, by proxy, how the brain response is modulated by sentence-level prosody, lexical semantics, and compositional structure and meaning. Our findings show that 1) the neuronal response is driven by compositional structure and meaning, beyond both acoustic-prosodic and lexical information; and 2) the brain most closely tracks the most structured representations on each of the distinct timescales we analyzed. To our knowledge, this is the first study to systematically disentangled the contribution of linguistic content from its timing and rhythm; the existing MI effects in the literature could have been driven by rhythmicity in the stimulus simply because no linguistically-informed controls had been tested. Additionally, our data demonstrates cortical tracking of language without a non-linguistic task (viz., syllable counting and outlier trial detection for Ding et al., and target-detection tasks for most other MI studies of speech processing). We show that oscillatory activity attunes to structured and meaningful content at distinct timescales, suggesting that phase synchronization reflects computations related to inferring linguistic representations from speech, and not merely the tracking of rhythmicity or timing. We discuss these findings in more detail below.

Using Mutual Information analysis, we quantified the degree of speech tracking in frequency bands corresponding to the timescales at which linguistic structures (phrases, words, and syllables) could be inferred from our stimuli. On the phrase timescale, we found that sentences had the most shared information between stimulus and response. Crucially, this is not merely a chunking mechanism (e.g., Bonhage et al., 2017; Ghitza, 2017) – participants could have “chunked” the word lists (which have their own naturally produced non-sentential prosody) into units of adjacent words, and the jabberwocky items into prosodic units. Instead, we argue that, on the timescale of phrases, the dominating process appears to be processing of compositional semantic structure, above and beyond prosodic chunking and word-level meaning.

Note that we do not take this observed increase in MI for sentences compared to jabberwocky items and word lists as evidence for an intrinsic “phrase-“, or “sentence-level oscillator”. Rather, we interpret the graded differences in MI at this timescale as a manifestation of the cortical computations that may occur during language comprehension – here, we observe them in the delta frequency range because that is the timescale on which higher-level linguistic units occur in our stimuli. Crucially, our experiment shows that the observed phase alignment of neural responses to inferred linguistic structures is not simply driven by the (quasi-)periodic occurrence of these structures at a given frequency: Both sentences and jabberwocky stimuli contained periodically occurring “phrases”, but our results show that the brain aligns more to those periodically occurring units when they contain meaningful information and are thus relevant for linguistic processing. These results are further corroborated by an additional, exploratory analysis using abstracted linguistic annotations of the stimuli (cf. Brodbeck et al., 2018).

On the word timescale, we found significantly higher MI between stimulus and brain response for sentences compared to jabberwocky items. We tentatively take this finding to indicate that, at the word timescale, the dominant process appears to be context-dependent word recognition – perhaps based in perceptual inference. Note, however, that we also found enhanced MI on the word timescale between stimuli and brain response for word lists compared to jabberwocky items when computing MI over all electrodes. Here, listeners could not have processed words within the context of phrases or sentences, which makes it somewhat difficult to integrate these results.

Many previous studies have shown that attention can modulate neural entrainment (e.g., Calderone et al., 2014; Ding & Simon, 2012; Golumbic et al., 2013; Haegens et al., 2011; Lakatos et al., 2013). Importantly, Ding et al. (2018) found that attention to a speech stimulus is required in order for neural tracking beyond the syllable envelope to be observed, suggesting that combining syllables into words and phrases requires attention. In our current experiment, participants were instructed to attentively listen to the audio recordings in all conditions, but it is possible that “attending to sentences” might be easier than “attending to jabberwocky items”, and that listeners are generally more likely to pay attention to higher-level structures in intelligible and meaningful speech. As such, we cannot rule out the possibility that our effects might be influenced by a mechanism that is based on attentional control. Note, however, that it is very difficult to disentangle “attention” from “comprehension” in this kind of argument – meaningful information within a stimulus can arguably only lead to increased attention if it is comprehensible. We plan to investigate these questions in more detail in future experiments.

Overall, the pattern of results is consistent with cue-integration-based models of language processing (Martin, 2016, 2020), where the activation profile of different populations of neurons over time is what encodes the linguistic structure as it is inferred from sensory correlates in real-time (Martin & Doumas, 2017). Martin’s (2016, 2020) model of language processing builds on and extends neurophysiological models of cue integration (e.g., Ernst & Bülthoff, 2004; Fetsch, DeAngelis, & Angelaki, 2013; Landy, Banks, & Knill, 2011; see Toscano & McMurray, 2015; and Norris & McQueen, 2008, for cue-integration-based models of speech and word recognition). The underlying mechanism of cue integration relies on only two core computations: summation and normalization, both of which have been proposed as canonical neural computations (e.g., Carandini & Heeger, 1994, 2012). Multiple cues (which can, in principle, be any piece of sensory information that is available in a given situation) are combined via summation and integrated via normalization in order to arrive at a robust percept (e.g., Ernst & Bülthoff, 2004). Cues are associated with corresponding weights, which can be dynamically updated in order to account for the fact that not all cues are equally reliable (or even available) in any given situation (see also Bates & MacWhinney, 1989, and Bornkessel & Bornkessel-Schlesewsky, 2006, for models of sentence processing that posit competition between different linguistic percepts as a result of cue validity and ranking). As such, the process of integrating multiple cues into a percept can be thought of as an inference problem (e.g., Landy et al., 2011). Martin (2016, 2020) proposed that, during all stages of language processing, the brain might draw on these same neurophysiological computations. Crucially, inferring linguistic representations from speech sounds requires not only bottom-up, sensory information, but also top-down, memory-based cues (e.g., Marslen-Wilson, 1987; Martin, 2016; Kaufeld et al., 2019). Martin (2016, 2020) therefore suggested that cue integration during language comprehension is an iterative process, where cues that have been inferred from the acoustic signal can, in turn, become cues for higher levels of processing. The pattern of findings in our current experiment strongly speaks to cue-integration-based models of language comprehension: We observe that neuronal tracking of the speech signal is enhanced for timescales on which meaningful linguistic units can be inferred, suggesting that phase alignment of populations of neurons might, indeed, encode the generation of inference-based linguistic representations (Martin & Doumas, 2017).

We found that phase alignment was significantly enhanced for higher-level linguistic structures, such as phrases, as opposed to syllables. These results are in line with previous findings from Keitel et al. (2018; see also Gross et al., 2013), who reported higher degrees of MI in the delta compared to the theta frequency band, corresponding to the frequency in which the most structured representations occurred in their stimuli. However, because we observed the same pattern of effects for forward *and* backward conditions, our data cannot speak to *why* tracking is enhanced at lower frequencies, and whether this is mainly due to confounds with 1/f brain activity (cf. Ouyang et al., 2019). Exploring in more detail why MI between brain and acoustic signal appears to be consistently highest at lower frequencies will be an interesting avenue for future research. To that end, we believe that studying naturalistic stimuli will further enhance our understanding of spoken language comprehension (Alday, 2019; see also Alexandrou et al., 2018).

There are, of course, many other open questions that arise from our results. Perhaps most obviously (although presumably limited by the resolution of time-frequency analysis), it would be interesting to investigate how “far” cue integration can be traced during more natural language comprehension situations (cf. Alday, 2019, Alexandrou et al., 2018). To what degree are higher-level linguistic cues, such as sentential, referential, contextual, or pragmatic information, encoded in the neural response? Another interesting avenue for future research would be to investigate whether similar patterns can be observed during language production. Martin (2016, 2020) suggested that not only language comprehension, but also language production draws on principles of cue integration. Observing similar oscillatory patterns during language production experiments, where speakers are actively generating linguistic units, would yield strong supporting evidence for cue-integration-based accounts of language processing. Finally – and consequentially, if cue integration underlies both comprehension and production processes –, we would be curious to learn more about cue integration “in action”, specifically during dialogue settings, where interlocutors comprehend and plan utterances nearly simultaneously.

In summary, this study showed that speech tracking is sensitive to both linguistic structure and meaning, above and beyond the frequency at which they occur in the stimulus. In other words: Content determines tracking, not just timescale. This extends previous findings and advances our understanding of spoken language comprehension in general, because our experimental manipulation allows us, for the first time, to disentangle the influence of linguistic structure and meaning on the neural response from the timescales at which that linguistic information occurs in the input. Cue-integration-based models of language processing (Martin, 2016, 2020; Martin & Doumas, 2017) offer a neurophysiologically plausible, mechanistic explanation for our results.

## Acknowledgments

We would like to thank Anne Keitel for sharing her expertise and scripts for the Mutual Information analysis, Laurel Brehm for statistical advice, Joe Rodd and Merel Maslowski for help with the experiment visualization (Figure 1), Annelies van Wijngaarden for lending her voice for stimulus recording, Karthikeya Kaushik, Zina al-Jibouri and Dylan Opdam for research assistance, and Micha Heilbron for feedback on an earlier draft of this manuscript. AEM was supported by the Max Planck Research Group “Language and Computation in Neural Systems” and by the Netherlands Organization for Scientific Research (grant 016.Vidi.188.029).

## Supplemental Materia1

### Stimuli

1. *Sentence*: Vlotte meesters schenken wijsheid en een aardig kindje schildert sterren. *(Chatty teachers offer wisdom and a nice child paints stars.)* *Jabberwocky*: Snatte waasters scharken wielheid en een aallig wundje schurdert sperben. *Word list*: schildert schenken vlotte aardig wijsheid kindje meesters sterren en een
2. Gekke meisjes snijden uien en de scherpe messen maken wondjes. *(Crazy girls cut onions and the sharp knives cause wounds.)* Gelpe muikjes floeden euer en de strerbe letsen lapen wouwses. uien wondjes meisjes messen de en snijden maken gekke scherpe
3. Kleine obers tapten biertjes en de domme gasten breken borden. *(Little waiter tapped beers and the stupid guests break plates.)* Spiene abels pipten beeltjes en de lolme gonten flepen varden. tapten breken kleine domme de en borden obers biertjes gasten
4. Lange mannen bouwen huisjes en de lieve honden brengen planken. *(Tall men build houses and the sweet dogs bring boards.)* Lalve wanzen botren raasjes en de reeve rorden brargen sponken. planken mannen huisjes honden de en bouwen brengen lange lieve
5. Trotse moeders hebben baby’s en de lieve oma’s geven snoepjes. *(Proud mothers have babies and the sweet grandmas give sweets.)* Pletse hijders rabben obis en de rieze bawun beben vliepjes. hebben geven trotse lieve de en snoepjes moeders baby’s oma’s
6. Goede sporters renden rondjes en de grote wolken bieden schaduw. *(Good athletes ran laps and the big clouds provide shadow.)* Vijde spenters rarden rouwses en de spode delken vuiden scharub. schaduw sporters rondjes wolken de en renden bieden goede grote
7. Stoute muizen knagen gaten en de boze huurders haten dieren. *(Naughty mice gnaw holes and the angry tenants hate animals.)* Stemte mieven snamen vaden en de vone hinkders doten weiren. knagen haten stoute boze de en gaten muizen huurders dieren
8. Slimme eekhoorns vinden nootjes en de groene kikkers vangen vliegjes. *(Smart squirrels find nuts and the green frogs catch flies.)* Plemme oekboorns ganden zietjes en de broeze wokkers gongen snoegjes. eekhoorns nootjes kikkers vliegjes de en vinden vangen slimme groene
9. Bange ridders zoeken toevlucht en de gouden sleutel opent deuren. *Frightened knights seek refuge and the golden key opens doors.)* Garge ludders nijken toepricht en de gatden speetel ogens weiren. zoeken opent bange gouden de en sleutel ridders toevlucht deuren
10. Blauwe visjes zwemmen baantjes en de grijze kippen horen piepjes. *(Blue fish swim laps and the grey chickens hear beeps.)* Braube bispes knimmen gaantres en de brijne dappen lolen peugjes. kippen visjes baantjes piepjes de en zwemmen horen blauwe grijze
11. Grote leeuwen vinden voedsel en de jonge schapen blijken geitjes. *(Big lions find food and the young sheep turn out to be goats.)* Spode loorden ginten baadsel en de jarge straben ploeken gaukjes. vinden lijken grote jonge de en voedsel leeuwen schapen geitjes
12. Kwade jongens breken glazen en de strenge juffen schrijven regels. *(Evil boys break glasses and the strict teachers write rules.)* Smate jargens drepen flaven en de strelle ceffen schroezen lemels. jongens juffen glazen regels de en breken schrijven kwade strenge
13. Zieke kindjes krijgen appels en de kalme zusters breien sokken. *(Sick children get apples and the calm sisters knit socks.)* Neike wundjes spijmen atsels en de malge nutters pleuen senken. krijgen breien zieke kalme de en kindjes appels zusters sokken
14. Warme landjes hebben strandjes en de korte dagen brengen vreugde. *(Warm lands have beaches and the short days bring joy.)* landjes strandjes dagen vreugde de en hebben brengen warme korte Marle lerkjes mobben strastpes en de warte lapen spelgen fleufde.
15. Zwarte geiten proefden suiker en de rotte tanden hebben gaten. *(Black goats taste sugar and the rotten teeth have holes.)* Flakte beuten praasden feeker en de hatte palden mabben voten. proefden hebben zwarte rotte de en gaten geiten suiker tanden
16. Rode mieren dragen takken en de wilde katten vangen vogels. *(Red ants carry branches and the wild cats catch birds.)* Lote keeren tramen tenken en de kelde lutten gargen valmen. takken mieren katten vogels de en dragen vangen rode wilde
17. Grauwe wolken brengen regen en de zware buien breken takken. *(Grey clouds bring rain and the heavy showers break branches.)* Kraube louken pletgen lepen en de plave gijen smesen tonken. brengen breken grauwe zware de en buien wolken regen takken
18. Blije artsen helpen mensen en de oude tantes hebben nichtjes. *(Happy doctors help people and the old aunts have nieces.)* Ploeie alfjen hospen miksen en de aide paltes labben zechtjes. artsen mensen tantes nichtjes de en helpen hebben oude blije
19. Houten tafels hebben laatjes en de ronde knikkers lijken druiven. *(Wooden tables have drawers and the round marbles look like grapes.)* Hemten pacels libben raakjes en de dande vlokkers woeken driezen. lijken houten hebben ronde de en tafels laatjes knikkers druiven
20. Zwarte laarzen trekken aandacht en de vreemde mannen schrobben vloeren. *Black boots attract attention and the strange men are scrubbing floors.)* Knorte raarnen grikken aangucht en de smijmde lonnen schrimben knijren. vloeren laarzen aandacht mannen de en trekken schrobben zwarte vreemde
21. Snelle jagers volgen spoortjes en de tamme hazen leggen keutels. *(Fast hunters follow tracks and the tame hares lay turds.)* Flolle cavers valmen vleertjes en de torme lamen lelmen weitels. volgen leggen snelle tamme de en jagers spoortjes keutels hazen
22. Leuke otters zoeken visjes en de grote leeuwen bijten mensen. *(Nice otters look for fish and the big lions bite people.)* Rauke akters nijken vaspes en de plode loorten gijden molsen leeuwen otters visjes mensen de en zoeken bijten leuke grote
23. Stille meisjes mengen sapjes en de rijke zeilers slurpen koffie. *(Silent girls mix juices and the rich sailors sip coffee.)* Stimpe muikjes lelgen sekjes en de lijse neulers plunpen wiffie. mengen slurpen stille rijke de en zeilers meisjes sapjes koffie
24. Trotse slagers snijden biefstuk en de blije klanten kopen worstjes. *(Proud butchers cut beef and the happy customers buy sausages.)* Bletse tramers proeden vaafstuk en de knoeie sponten women wuchtjes. biefstuk worstjes klanten slagers de en snijden kopen trotse blije
25. Knappe schilders winnen prijsjes en de wilde paarden aten peren. *(Handsome painters win prizes and the wild horses eat pears.)* Flippe scharders dinzen proekjes en de kelde deurden usen remen. winnen aten knappe wilde de en schilders prijsjes paarden peren
26. Stompe messen snijden broodjes en de paarse pennen schrijven woorden. *(Blunt knives cut sandwiches and the purple pens write words.)* Starpe retsen knoeden braaljes en de waadse parnen schroezen moorten. messen broodjes pennen woorden de en snijden schrijven stompe paarse
27. Bruine apen zoeken vruchten en de witte schapen aten blaadjes. *(Brown monkeys seek fruit and the white sheep eat leaves.)* Driene onen nijken smechten en de kette straven oken bleegjes. zoeken aten bruine witte de en apen schapen blaadjes vruchten
28. Oude opa’s snoeien heggen en de lieve oma’s bakten koekjes. *(Old grandpas prune hedges and the sweet grandmas baked cookies.)* Adde obos knooien relgen en de ruive onis borten moefjes. koekjes opa’s heggen oma’s de en snoeien bakten oude lieve
29. Trieste zwemmers schrijven brieven en de lompe zangers zingen liedjes. *(Sad swimmers write letters and the rude singers sing songs.)* Breeste knimmers schroezen pleiven en de laspe zallers zannen riefjes. schrijven zingen trieste lompe de en brieven liedjes zangers zwemmers
30. Stoere vaders prikken gaten en de vlotte moeders koken uien. *(Tough fathers poke holes and the chatty mothers cook onions.)* Stijne gaters drekken vaden en de knette hijders mosen auer. moeders vaders gaten uien de en prikken koken stoere vlotte
31. Mooie vogels zingen wijsjes en de gekke meiden wassen kleren. *(Beautiful birds sing tunes and the crazy girls wash clothes.)* Woeie govels zanpen waadjes en de gesse kieden pansen pleven. zingen wassen mooie gekke de en vogels wijsjes kleren meiden
32. Knappe zangers geven feestjes en de stoere werklui kopen biertjes. *(Handsome singers give parties and the tough workers buy beers.)* Smippe zalpers beben feursjes en de steepe werfmui rogen booltjes. zangers werklui biertjes feestjes de en geven kopen knappe stoere
33. Vieze kwallen prikken duikers en de dunne vissers huurden bootjes. *(Dirty jellyfish sting divers and the thin fishermen rented boats.)* Beeze flollen kwokken keekers en de murne bitsers lutsden boepjes. prikken huurden vieze dunne de en duikers vissers bootjes kwallen
34. Sterke vaders dragen dochters en de mooie meisjes zoenden jongens. *(Strong fathers carry daughters and the beautiful girls kissed boys.)* Sperre goders tramen wichters en de moene miekjes nienden jorlens. dochters meisjes vaders jongens de en dragen zoenden sterke mooie
35. Lompe kappers knippen haren en de vlugge klussers bouwen muren. *(Rude hairdressers cut hair and the quick handymen built walls.)* Lolle mippers vrappen lalen en de snigge spessers botren luven. knippen bouwen lompe vlugge de en muren kappers klussers haren
36. Leuke vrouwen spelen cello en de dikke drummers poetsten trommels. *(Nice women play cello and the fat drummers cleaned drums.)* Mauke smouven pleren jeldo en de wokke plurmers peursten spolmels. vrouwen drummers cello trommels de en spelen poetsten leuke dikke
37. Arme vrouwen poetsten schoenen en de trage laptops brengen spanning. *(Poor women polished shoes and the slow laptops cause tension.)* Orle vrulwen poonsten scheemen en de drame lanteps spelgen klanzing. poetsten brengen arme trage de en vrouwen spanning schoenen laptops
38. Lieve meisjes plukten appels en de schuwe jongens vrezen hoogtes. *(Sweet girls picked apples and the shy boys are afraid of heights.)* Reeve muipjes slunten atjels en de schine jargens flenen haaites. meisjes hoogtes appels jongens de en plukten vrezen lieve schuwe
39. Drukke winkels lokten klanten en de lange mannen kopen schoenen. *(Busy stores attracted customers and the tall men buy shoes.)* Spunke linsels lurten spalten en de lalve wanzen loben scheegen. lokten kopen drukke lange de en winkels klanten schoenen mannen
40. Gekke jongens pesten eenden en de zieke meiden poetsten tanden. *(Crazy boys harrass ducks and the sick girls brushed their teeth.)* Gelse jormens tetten oelden en de neike kieden peugsten palden. tanden jongens meiden eenden de en pesten poetsten gekke zieke
41. Vreemde vrouwen hebben heimwee en de trouwe buren zenden brieven. *(Strange women are homesick and the loyal neighbors send letters.)* Smuimde flouven rabben heipwij en de blouve guven zarden pleiven. hebben zenden vreemde trouwe de en heimwee vrouwen brieven buren
42. Kleine baby’s horen liedjes en de wijze kerels lezen kranten. *(Small babies hear songs and the wise guys read newspapers.)* Speune bawus ronen riefjes en de moeze lenels remen sponten. baby’s kerels liedjes kranten de en horen lezen kleine wijze
43. Saaie buren kopen borden en de jonge kindjes pakten snoepjes. *(Boring neighbors buy plates and the young children grabbed sweets.)* Siere gulen loben girden en de jelge wirtjes penten vloesjes. pakten saaie kopen jonge de en buren borden kindjes snoepjes
44. Snelle schaatsers vinden gaatjes en de kleine jongens spelen voetbal. *(Fast skaters find holes and the little boys play soccer.)* Flolle schijnsers ginten geekjes en de speene jargens sleren boenbel. schaatsers voetbal gaatjes jongens de en vinden spelen snelle kleine
45. Witte paarden trekken koetsen en de saaie vorsten wenkten burgers. *(White horses pull carriages and the boring monarchs greet civilians.)* Kette peenden drakken dietsen en de soene viksten lankten vurmers. trekken wenkten witte saaie de en paarden vorsten burgers koetsen
46. Stille schilders belden vrienden en de roze scooter levert pizza. *(Quiet painters called friends and the pink scooter delivers pizza.)* Stimpe schadders benten fleunden en de lone sjaater rezert pixta. schilders scooter vrienden pizza de en roze stille levert belden
47. Oude mensen rijden bussen en de toffe oma’s maken grapjes. *(Old people ride busses and the cool grandmas make jokes.)* Amde molsen mieden gossen en de puffe onos lapen spipjes. rijden maken oude toffe de en bussen grapjes oma’s mensen
48. Brede tantes zoenden wangen en de malle neven roken jointjes. *(Sturdy aunts kissed cheeks and the silly nephews smoked joints.)* Krete pastes noonden largen en de kolle zeben kosen jarstjes. wangen neven jointjes tantes de en zoenden roken brede malle
49. Jonge bakkers maken broden en de nette klanten ruiken taartjes. *(Young bakers make bread and the neat customers smell cake.)* Jorle bassers hapen smoten en de zutte spalten hieken toordjes. maken ruiken jonge nette de en broden taartjes klanten bakkers
50. Klamme handen voelen muren en de knusse kachels drogen kleren. *(Damp hands feel walls and the cozy ovens dry clothes.)* Klorme londen vijren luven en de vresse maspels blomen spemen. handen muren kachels kleren de en voelen drogen klamme knusse
51. Snelle zwemmers slurpen ijsthee en de vieze kwallen zoeken water. *(Fast swimmers slurp iced tea and the dirty jellyfish look for water.)* Knille knummers plunpen ijfkwee en de bieve smollen nijken lates. slurpen zoeken vieze snelle de en zwemmers ijsthee kwallen water
52. Bange helden plukken bloemen en de bruine vogels halen takken. *(Frightened heroes pick flowers and the brown birds fetch branches.)* Garge ralden spunken drijmen en de druize gomels paven mukken. helden bloemen vogels takken de en plukken halen bange bruine
53. Vlotte otters bouwen dammen en de snelle hazen doden kevers. *(Smooth otters build dams and the fast hares kill beetles.)* Zwitte olders botren lemmen en de vralle lamen zoten mezers. bouwen doden vlotte snelle de en otters dammen hazen kevers
54. Saaie meesters geven lessen en de vele brieven worden stapels. *(Boring teachers give lessons and the many letters become piles.)* Soene waasters beben hussen en de bene pleeven rarden stagelt. meesters lessen stapels brieven de en geven worden saaie vele
55. Luikse wafels stillen honger en de rotte appels krijgen schimmel. *(Liège waffles satisfy hunger and the rotten apples are getting moldy.)* Ruipre lafelt stimpen morger en de hatte ampelt spijmen schurmel. krijgen stillen luikse rotte de en wafels honger appels schimmel
56. Enge slangen eten muizen en de grote kippen leggen eitjes. *(Scary snakes eat mice and the big chickens lay eggs.)* Elme spalgen eber mienen en de vlode dappen relgen eutres. slangen muizen kippen eitjes de en eten leggen enge grote
57. Scherpe scharen knippen blaadjes en de snelle auto’s rijden rondjes. *(Sharp scissors cut leaves and the fast cars drive laps.)* Strerbe stranen smeppen bleegjes en de flalle euvos loeden lortjes. knippen rijden scherpe snelle de en scharen blaadjes auto’s rondjes
58. Luie tieners dekken tafels en de dikke dames brengen koffie. *(Lazy teenagers set tables and the fat ladies bring coffee.)* Reie teezers lenken mabels en de wokke lapes brargen wiffie. tieners tafels dames koffie de en dekken brengen luie dikke
59. Zure bessen maken vlekken en de zachte bijen maken honing. *(Sour berries make stains and the soft bees make honey.)* Nume betjen lasen zwokken en de nochte guien lapen moping. maken maken zure zachte de en bessen bijen vlekken honing
60. Vieze mannen smeren olie en de toffe ouders kopen kaartjes. *(Dirty men smear oil and the cool parents buy tickets.)* Bieve wanzen fleven oree en de piffe amders homen koostjes. mannen olie ouders kaartjes de en smeren kopen vieze toffe
61. Franse schilders verven muren en de Vlaamse bakkers kneden brooddeeg. *(French painters paint walls and the Flemish bakers were kneading bread dough.)* Flanje schunders bernen lunen en de knuimse garkers dreten slooddieg. verven kneden Franse Vlaamse de en schilders muren bakkers brooddeeg
62. Vlotte lopers maken meters en de boze tieners slopen ruiten. *(Fast runners make meters and the angry teenagers demolish windows.)* Snette rogeres dapen peders en de vone teezers spomen hieten. lopers meters tieners ruiten de en maken slopen vlotte boze
63. Gele bloemen lokken bijen en de brakke mensen drinken water. *(Yellow flowers attract bees and the hungover people drink water.)* Bese drijmen lunken veien en de plokke moksen plinsen rates. drinken lokken gele brakke de en bloemen bijen mensen water
64. Zware tassen breken ruggen en de natte sokken brengen blaren. *(Heavy bags break backs and the wet socks cause blisters.)* Plane pansen flesen ruflen en de zaste sunken spelgen spalen. tassen ruggen sokken blaren de en breken brengen zware natte
65. Losse spijkers sieren muren en de bange muizen graven hollen. *(Loose nails adorn walls and the scared mice dig holes.)* Lunse kloekers suinen luven en de garge mienen slazen mellen. sieren graven losse bange de en spijkers muren muizen hollen
66. Dronken rappers lopen blauwtjes en de bolle katten aten brokjes. *(Drunk rappers are turned down and the fluffy cats ate kibble.)* Plorken lippers women bleuntjes en de galle mitten uken slekjes. lopen aten dronken bolle de en rappers blauwtjes katten brokjes
67. Knappe ridders redden levens en de grote paarden winnen prijsjes. *(Handsome knights save lives and the big horses win prizes.)* Flippe ludders hefden rezens en de spode deurden dinzen proekjes. ridders paarden prijsjes levens de en redden winnen knappe grote
68. Kille zussen stelen spullen en de woeste ouders straffen broertjes. *(Cold sisters steal things and the savage parents punish brothers.)* Wulle zetsen speven krillen en de deuste agders stropfen brooltjes. stelen straffen kille woeste de en zussen spullen ouders kindjes
69. Slome slakken aten sprietjes en de valse hommels steken kindjes. *(Slow snails ate grass blades and the false bumblebees sting children.)* Ploge spikken osen sproekjes en de vikse lompels speven wirtjes. slakken sprietjes hommels kindjes de en aten steken slome valse
70. Rotte appels brengen ziektes en de gulle fietsers zingen liedjes. *(Rotten apples bring diseases and the generous cyclists sing songs.)* Hatte ammels spelgen nientes en de gumpe feunsers zansen riegjes. brengen zingen rotte gulle de en appels ziektes fietsers liedjes
71. Gekke buren maken worstjes en de dikke neven snoepen taarten. *(Crazy neighbors make sausages and the heavy cousins snack on cake.)* Gelse gulen dapen wuchtjes en de mekke zeben smijpen tijnten. snoepen maken gekke dikke de en buren worstjes neven taarten
72. Dunne meisjes drinken sapjes en de witte duiven aten bonen. *(Thin girls drink juices and the white pigeons ate beans.)* Durre muipjes plinsen sutjes en de kette wuizen uken goven. meisjes sapjes duiven bonen de en drinken aten witte dunne
73. Knappe mannen strikken veters en de kleine muizen horen piepjes. *(Handsome men tie laces and the little mice heard beeps.)* Flippe wanzen strissen getels en de spiene mieven rolen peugjes. strikken horen knappe kleine de en mannen muizen veters piepjes
74. Vieze messen snijden appels en de snelle jongens gooien ballen. *(Dirty knives cut apples and the fast guys throw balls.)* Beize letsen floeden ammels en de flolle jargens goenen garlen. messen appels jongens ballen de en snijden gooien vieze snelle
75. Trage mensen kopen broden en de mooie vrouwen bakken taarten. *(Slow people buy bread and the beautiful women baked pies.)* Klame helsen lomen droten en de moere smouven galken peurten. kopen bakken trage mooie de en mensen broden vrouwen taarten
76. Luie honden ruiken voedsel en de warme broodjes hebben pitjes. *(Lazy dogs smell food and the hot sandwiches contain seeds.)* Reue rorden hieken baadsel en de marle braagjes rabben pekjes. voedsel honden broodjes pitjes de en ruiken hebben luie warme
77. Boze ouders geven straffen en de rode vossen graven kuilen. *(Angry parents give punishments and the red foxes dig pits.)* Vome adkers besen strinfen en de lote gussen brazen mielen. geven graven boze rode de en ouders vossen kuilen straffen
78. Lieve meiden schrijven boeken en de lichte kamers hebben ramen. *(Sweet girls write books and the bright rooms have windows.)* Reive kieden schroezen vijken en de rachte lapers mabben dapen. meiden boeken kamers ramen de en schrijven hebben lieve lichte
79. Gouden munten hebben waarde en de vreemde vogels fluiten liedjes. *(Golden coins have value and the strange birds whistle songs.)* Gudden kurten rabben laalde en de floemde govels vrieten leugjes. hebben fluiten gouden vreemde de en munten waarde vogels liedjes
80. Slome treinen hebben stoelen en de rijke boeren voeden koeien. *(Slow trains have seats and the rich farmers feed cows.)* treinen stoelen boeren koeien de en hebben voeden slome rijke Ploge pleenen labben stijren en de loeke goelen vuiten woenen.

### Supplementary Figures

**Supplementary Figure A.**
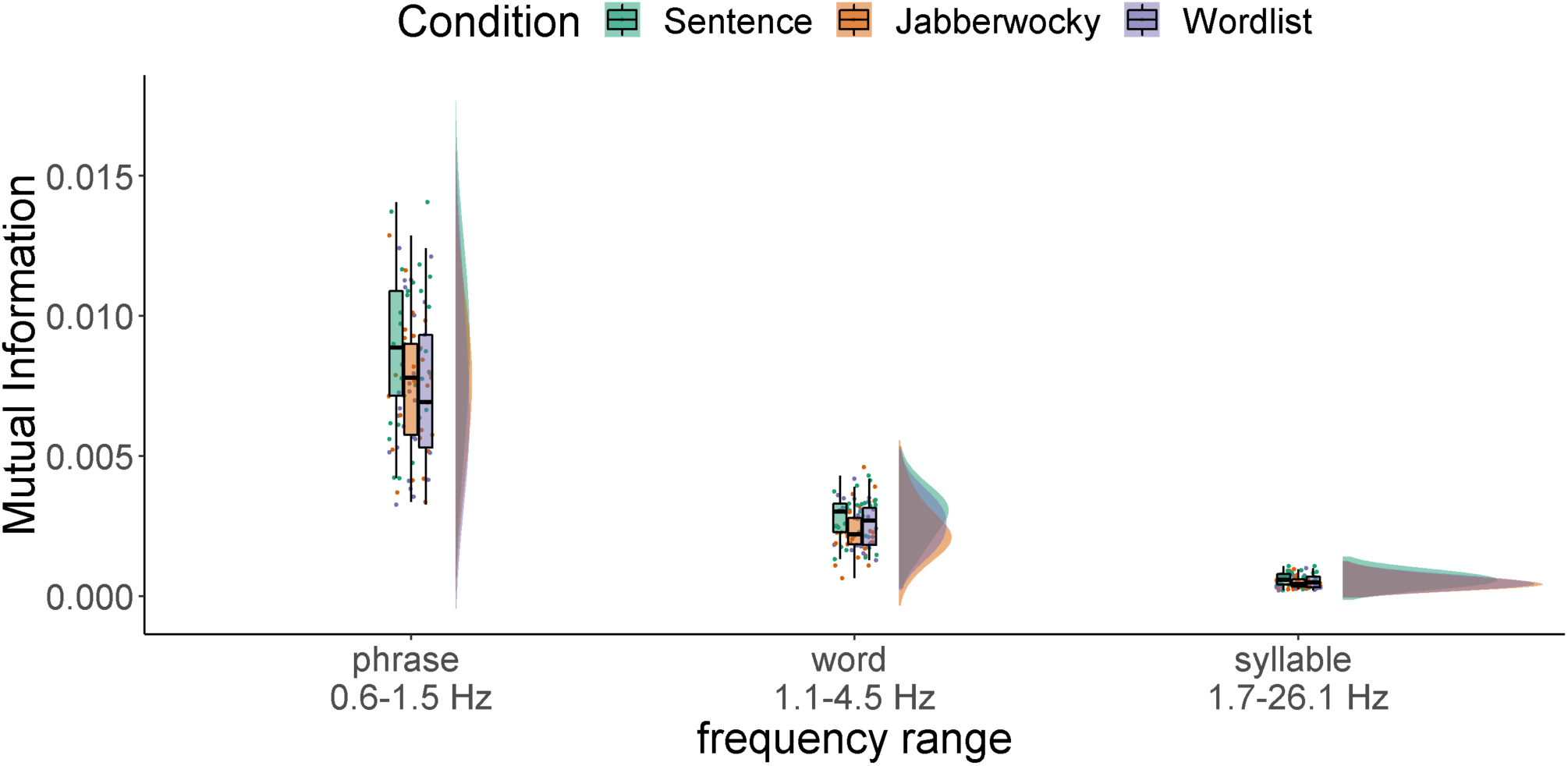
Phase MI between speech signal and brain response for wider frequency bins. Frequency bins were calculated based on maximum and minimum durations (across all stimuli and conditions) for phrases, words, and syllables (phrases: 0.6-1.5 Hz; words: 1.1-4.5 Hz; syllables: 1.7-26.1 Hz). Note that bins overlap because, naturalistically, durations of linguistic units overlapped. MI was calculated over the entire head. Differences between Sentence – Jabberwocky and Sentence – Wordlist were significant on the phrase and syllable timescale; between Sentence – Wordlist on the word timescale. No significant differences were found for the backward conditions.

**Supplementary Figure B.**
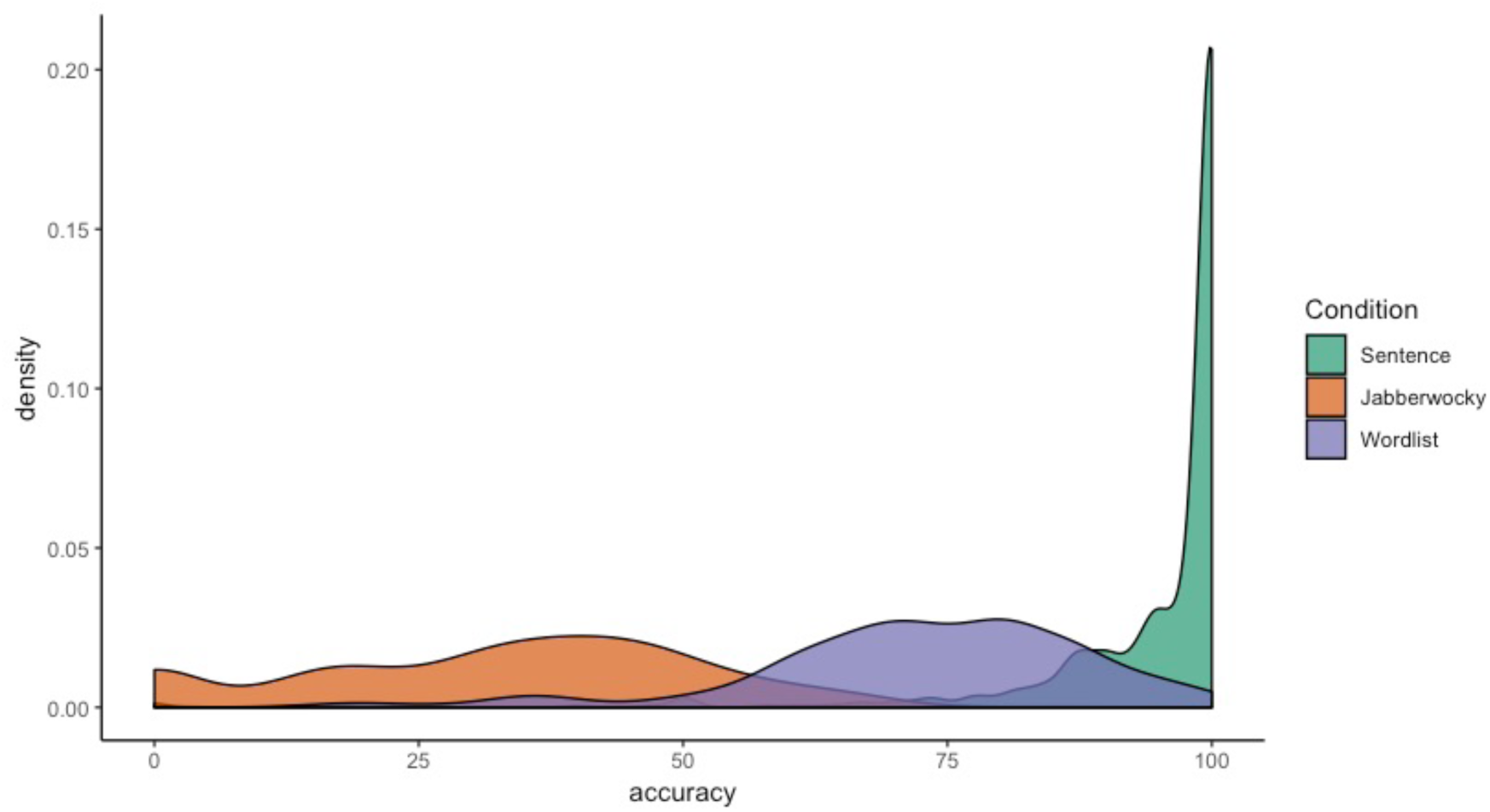
Density distribution of accuracy scores (in %; calculated as string token sort ratio between excepted and reported responses during the pretest) for Sentence (green), Jabberwocky (orange) and Wordlist (purple) items.

## Footnotes

1 That is, even though phrases, clauses, and sentences can at times be cued by acoustic-prosodic means (e.g., silent pauses, filled pauses, final lengthening, fundamental frequency reset, etc.; cf. Bosker, Pinget, Quené, Sanders, & De Jong 2013), they are less easily detectable in the acoustic signal compared to syllables (which are obvious from a clear peak in the modulation spectrum of speech; Ding et al., 2014, 2016).

2 Note that the problem the brain faces during spoken language comprehension is even more complex than this, because the timescales of linguistic units can highly overlap, even within a single sentence. Populations of neurons that “entrain” to words will thus also have to be sensitive to information that occurs outside of these – rather narrow – frequency bands. We also computed MI values within wider frequency bands that were based on the absolute minimum and maximum duration of units at each timescale across all stimuli (phrases: 0.6-1.5 Hz; words: 1.1-4.5 Hz; syllables: 1.7-26.1 Hz), see Figure A (Supplemental Materials) for a visualization of results.

3 We thank an anonymous reviewer for inspiring this exploratory analysis.

## References

Alday, Phillip M. (2019). M/EEG analysis of naturalistic stories: a review from speech to language processing. Language, Cognition and Neuroscience, 34(4), 457–473.

Alexandrou, A. M., Saarinen, T., Kujala, J., & Salmelin, R. (2018). Cortical entrainment: what we can learn from studying naturalistic speech perception. Language, Cognition and Neuroscience, 1-13.

Allen, M., Poggiali, D., Whitaker, K., Marshall, T. R., & Kievit, R. A. (2019). Raincloud plots: a multi-platform tool for robust data visualization. Wellcome open research, 4.

Audacity Team (2014). Audacity(R): Free Audio Editor and Recorder.

Bates, E., & MacWhinney, B. (1989). Functionalism and the competition model. The crosslinguistic study of sentence processing, 3, 73–112.

Bates, D., Maechler, M., Bolker, B., & Walker, S. (2015). Fitting linear mixed-effects models using lme4. Journal of Statistical Software, 67(1), 1–48. doi: 10.18637/jss.v067.i01

Bever, T. G., & Poeppel, D. (2010). Analysis by Synthesis: A (Re-)Emerging Program of Research for Language and Vision. Biolinguistics 4(2-3), 174–200.

Boersma, Paul & Weenink, David (2017). Praat: doing phonetics by computer (Version 6.0.36).

Bonhage, C. E., Meyer, L., Gruber, T., Friederici, A. D., & Mueller, J. L. (2017). Oscillatory EEG dynamics underlying automatic chunking during sentence processing. NeuroImage, 152, 647–657. https://doi.org/10.1016/j.neuroimage.2017.03.018

Bosker, H. R., & Cooke, M. (2018). Talkers produce more pronounced amplitude modulations when speaking in noise. The Journal of the Acoustical Society of America, 143(2), EL121-EL126.

Bosker, H. R., & Ghitza, O. (2018). Entrained theta oscillations guide perception of subsequent speech: behavioural evidence from rate normalisation. Language, Cognition and Neuroscience, 33(8), 955–967.

Bosker, H. R., Pinget, A. F., Quené, H., Sanders, T., & De Jong, N. H. (2013). What makes speech sound fluent? The contributions of pauses, speed and repairs. Language Testing, 30(2), 159–175.

Bornkessel, I., & Schlesewsky, M. (2006). The extended argument dependency model: a neurocognitive approach to sentence comprehension across languages. Psychological review, 113(4), 787.

Brodbeck, C., Presacco, A., & Simon, J. Z. (2018). Neural source dynamics of brain responses to continuous stimuli: Speech processing from acoustics to comprehension. NeuroImage, 172, 162–174.

Calderone, D. J., Lakatos, P., Butler, P. D., & Castellanos, F. X. (2014). Entrainment of neural oscillations as a modifiable substrate of attention. Trends in cognitive sciences, 18(6), 300–309.

Carandini, M., & Heeger, D. J. (1994). Summation and division by neurons in primate visual cortex. Science, 264(5163), 1333–1336. https://doi.org/10.1126/science.8191289

Carandini, M., & Heeger, D. J. (2012). Normalization as a canonical neural computation. Nature Reviews Neuroscience, 13(1), 51–62. https://doi.org/10.1038/nrn3136

Cogan, G. B., & Poeppel, D. (2011). A mutual information analysis of neural coding of speech by low-frequency MEG phase information. Journal of neurophysiology, 106(2), 554–563.

Ding, N., Melloni, L., Yang, A., Wang, Y., Zhang, W., & Poeppel, D. (2017). Characterizing Neural Entrainment to Hierarchical Linguistic Units using Electroencephalography (EEG). Frontiers in Human Neuroscience, 11. https://doi.org/10.3389/fnhum.2017.00481

Ding, N., Patel, A. D., Chen, L., Butler, H., Luo, C., & Poeppel, D. (2017b). Temporal modulations in speech and music. Neuroscience & Behavioural Reviews, 81, 181–187. https://doi.org/10.1016/j.neubiorev.2017.02.011

Ding, N., Pan, X., Luo, C., Su, N., Zhang, W., & Zhang, J. (2018). Attention is required for knowledge-based sequential grouping: insights from the integration of syllables into words. Journal of Neuroscience, 38(5), 1178–1188.

Ding, N., Melloni, L., Zhang, H., Tian, X., & Poeppel, D. (2016). Cortical tracking of hierarchical linguistic structures in connected speech. Nature Neuroscience, 19(1), 158–164. https://doi.org/10.1038/nn.4186

Ding, N., & Simon, J. Z. (2012). Emergence of neural encoding of auditory objects while listening to competing speakers. Proceedings of the National Academy of Sciences, 109(29), 11854–11859.

Doelling, K. B., Arnal, L. H., Ghitza, O., & Poeppel, D. (2014). Acoustic landmarks drive delta–theta oscillations to enable speech comprehension by facilitating perceptual parsing. Neuroimage, 85, 761–768.

Ernst, M. O., & Bülthoff, H. H. (2004). Merging the senses into a robust percept. Trends in Cognitive Sciences, 8(4), 162–169. https://doi.org/10.1016/j.tics.2004.02.002

Fetsch, C. R., DeAngelis, G. C., & Angelaki, D. E. (2013). Bridging the gap between theories of sensory cue integration and the physiology of multisensory neurons. Nature Reviews Neuroscience, 14(6), 429–442. https://doi.org/10.1038/nrn3503

Frank, S. L., & Yang, J. (2018). Lexical representation explains cortical entrainment during speech comprehension. PloS one, 13(5), e0197304.

Ghitza, O. (2017). Acoustic-driven delta rhythms as prosodic markers. Language, Cognition and Neuroscience, 32(5), 545–561. https://doi.org/10.1080/23273798.2016.1232419

Golumbic, E. M. Z., Ding, N., Bickel, S., Lakatos, P., Schevon, C. A., McKhann, G. M., … & Poeppel, D. (2013). Mechanisms underlying selective neuronal tracking of attended speech at a “cocktail party”. Neuron, 77(5), 980–991.

Gross, J., Hoogenboom, N., Thut, G., Schyns, P., Panzeri, S., Belin, P., & Garrod, S. (2013). Speech Rhythms and Multiplexed Oscillatory Sensory Coding in the Human Brain. PLoS Biology, 11(12), e1001752. https://doi.org/10.1371/journal.pbio.1001752

Haegens, S., & Golumbic, E. Z. (2018). Rhythmic facilitation of sensory processing: A critical review. Neuroscience & Biobehavioral Reviews, 86, 150–165.

Haegens, S., Händel, B. F., & Jensen, O. (2011). Top-down controlled alpha band activity in somatosensory areas determines behavioral performance in a discrimination task. Journal of Neuroscience, 31(14), 5197–5204.

Halle, M., & Stevens, K. N. (1962). Speech recognition: A model and a program for research. IRE Transactions of the PGIT, IT-8, 155–159.

Henry, M. J., Herrmann, B., & Obleser, J. (2014). Entrained neural oscillations in multiple frequency bands comodulate behavior. Proceedings of the National Academy of Sciences, 111(41), 14935–14940.

Henry, M. J., & Obleser, J. (2012). Frequency modulation entrains slow neural oscillations and optimizes human listening behavior. Proceedings of the National Academy of Sciences, 109(49), 20095–20100.

Howard, M. F., & Poeppel, D. (2010). Discrimination of speech stimuli based on neuronal response phase patterns depends on acoustics but not comprehension. Journal of neurophysiology, 104(5), 2500–2511.

Ince, R. A., Giordano, B. L., Kayser, C., Rousselet, G. A., Gross, J., & Schyns, P. G. (2017). A statistical framework for neuroimaging data analysis based on mutual information estimated via a gaussian copula. Human brain mapping, 38(3), 1541–1573.

Kaufeld, G., Ravenschlag, A., Meyer, A. S., Martin, A. E., & Bosker, H. R. (2019). Knowledge-based and signal-based cues are weighted flexibly during spoken language comprehension. Journal of Experimental Psychology: Learning, Memory, and Cognition.

Kayser, S. J., Ince, R. A. A., Gross, J., & Kayser, C. (2015). Irregular Speech Rate Dissociates Auditory Cortical Entrainment, Evoked Responses, and Frontal Alpha. Journal of Neuroscience, 35(44), 14691–14701. https://doi.org/10.1523/JNEUROSCI.2243-15.2015

Keitel, A., Gross, J., & Kayser, C. (2018). Perceptually relevant speech tracking in auditory and motor cortex reflects distinct linguistic features. PLOS Biology, 16(3), e2004473. https://doi.org/10.1371/journal.pbio.2004473

Keitel, A., Ince, R. A. A., Gross, J., & Kayser, C. (2017). Auditory cortical delta-entrainment interacts with oscillatory power in multiple fronto-parietal networks. NeuroImage, 147, 32–42. https://doi.org/10.1016/j.neuroimage.2016.11.062

Keuleers, E., & Brysbaert, M. (2010). Wuggy: A multilingual pseudoword generator. Behavior Research Methods 42(3), 627–633.

Kösem, A., Bosker, H. R., Takashima, A., Meyer, A., Jensen, O., & Hagoort, P. (2018). Neural entrainment determines the words we hear. Current Biology, 28(18), 2867–2875.

Kutas, M., & Federmeier, K. D. (2000). Electrophysiology reveals semantic memory use in language comprehension. Trends in cognitive sciences, 4(12), 463–470.

Kutas, M., Van Petten, C. K., & Kluender, R. (2006). Psycholinguistics electrified II (1994–2005). In Handbook of psycholinguistics (pp. 659-724). Academic Press.

Kuznetsova, A., Brockhoff, P. B., Christensen, R. H. B. (2017). lmerTest Package: Tests in Linear Mixed Effects Models. Journal of Statistical Software, 82(13), 1–26.

Lakatos, P., Musacchia, G., O’Connel, M. N., Falchier, A. Y., Javitt, D. C., & Schroeder, C. E. (2013). The spectrotemporal filter mechanism of auditory selective attention. Neuron, 77(4), 750–761.

Landy, M. S., Banks, M. S., & Knill, D. C. (2011). Ideal-Observer Models of Cue Integration. In J. Trommershäuser, K. Kording, & M. S. Landy (Eds.), Sensory Cue Integration (pp. 5–29). Oxford University Press. https://doi.org/10.1093/acprof:oso/9780195387247.003.0001

Length, R. (2018). emmeans: Estimated Marginal Means, aka Least-Square Means. R package version 1.3.0. https://CRAN.R-project.org/package=emmeans

Luo, H., & Poeppel, D. (2007). Phase patterns of neuronal responses reliably discriminate speech in human auditory cortex. Neuron, 54(6), 1001–1010.

Luo, H., & Poeppel, D. (2012). Cortical oscillations in auditory perception and speech: evidence for two temporal windows in human auditory cortex. Frontiers in psychology, 3, 170.

Luo, H., Liu, Z., & Poeppel, D. (2010). Auditory cortex tracks both auditory and visual stimulus dynamics using low-frequency neuronal phase modulation. PLoS biology, 8(8), e1000445.

Marslen-Wilson, W. D. (1987). Functional parallelism in spoken word-recognition. Cognition, 25(1–2), 71–102. https://doi.org/10.1016/0010-0277(87)90005-9

Martin, A. E. (2016). Language Processing as Cue Integration: Grounding the Psychology of Language in Perception and Neurophysiology. Frontiers in Psychology, 7. https://doi.org/10.3389/fpsyg.2016.00120

Martin, A. E. (2020). A compositional neural architecture for language. In press at Journal of Cognitive Neuroscience.

Martin, A. E., & Doumas, L. A. A. (2017). A mechanism for the cortical computation of hierarchical linguistic structure. PLOS Biology, 15(3), e2000663. https://doi.org/10.1371/journal.pbio.2000663

Meyer, L. (2018). The neural oscillations of speech processing and language comprehension: state of the art and emerging mechanisms. European Journal of Neuroscience, 48(7), 2609–2621.

Meyer, L., Henry, M. J., Gaston, P., Schmuck, N., & Friederici, A. D. (2016). Linguistic Bias Modulates Interpretation of Speech via Neural Delta-Band Oscillations. Cerebral Cortex. https://doi.org/10.1093/cercor/bhw228

Meyer, L., Sun, Y., & Martin, A. E. (2019). Synchronous, but not entrained: exogenous and endogenous cortical rhythms of speech and language processing. Language, Cognition and Neuroscience, 1-11.

Norris, D., & McQueen, J. M. (2008). Shortlist B: A Bayesian model of continuous speech recognition. Psychological Review, 115(2), 357–395. https://doi.org/10.1037/0033-295X.115.2.357

Obleser, J., & Kayser, C. (2019). Neural Entrainment and Attentional Selection in the Listening Brain. Trends in cognitive sciences.

Oostenveld, R., Fries, P., Maris, E., Schoffelen, JM (2011) FieldTrip: Open Source Software for Advanced Analysis of MEG, EEG, and Invasive Electrophysiological Data. Computational Intelligence and Neuroscience Volume 2011 (2011), Article ID 156869, doi:10.1155/2011/156869

Ouyang, G., Hildebrandt, A., Schmitz, F., & Herrmann, C. S. (2019). Decomposing alpha and 1/f brain activities reveals their differential associations with cognitive processing speed. NeuroImage, 116304.

Peelle, Jonathan E., & Davis, M. H. (2012). Neural Oscillations Carry Speech Rhythm through to Comprehension. Frontiers in Psychology, 3. https://doi.org/10.3389/fpsyg.2012.00320

Peelle, J. E., Gross, J., & Davis, M. H. (2013). Phase-Locked Responses to Speech in Human Auditory Cortex are Enhanced During Comprehension. Cerebral Cortex, 23(6), 1378–1387. https://doi.org/10.1093/cercor/bhs118

Poeppel, D., & Monahan, P. J. (2011). Feedforward and feedback in speech perception: Revisiting analysis by synthesis. Language and Cognitive Processes, 26(7), 935–951. https://doi.org/10.1080/01690965.2010.493301

R Core Team (2018). R: A language and environment for statistical computing. R Foundation for Statistical Computing, Vienna, Austria. https://www.R-project.org/

Schroeder, C. E., & Lakatos, P. (2009). Low-frequency neuronal oscillations as instruments of sensory selection. Trends in Neurosciences, 32(1), 9–18. https://doi.org/10.1016/j.tins.2008.09.012

Smith, Z. M., Delgutte, B., & Oxenham, A. J. (2002). Chimaeric sounds reveal dichotomies in auditory perception. Nature, 416(6876), 87.

Teng, X., Tian, X., Doelling, K., & Poeppel, D. (2018). Theta band oscillations reflect more than entrainment: behavioral and neural evidence demonstrates an active chunking process. European Journal of Neuroscience, 48(8), 2770–2782. https://doi.org/10.1111/ejn.13742

Toscano, J. C., & McMurray, B. (2015). The time-course of speaking rate compensation: effects of sentential rate and vowel length on voicing judgments. Language, Cognition and Neuroscience, 30(5), 529–543. https://doi.org/10.1080/23273798.2014.946427

Watanabe, H., Tanaka, H., Sakti, S., & Nakamura, S. (2019). Synchronization between overt speech envelope and EEG oscillations during imagined speech. Neuroscience research.

Zoefel, B., ten Oever, S., & Sack, A. T. (2018). The Involvement of Endogenous Neural Oscillations in the Processing of Rhythmic Input: More Than a Regular Repetition of Evoked Neural Responses. Frontiers in Neuroscience, 12. https://doi.org/10.3389/fnins.2018.00095

Zoefel, B., & VanRullen, R. (2015). The Role of High-Level Processes for Oscillatory Phase Entrainment to Speech Sound. Frontiers in Human Neuroscience, 9. https://doi.org/10.3389/fnhum.2015.00651

